# A strong-to-weak interaction shift during microbiome succession is coupled to colonizer-dependent antimicrobial resistance

**DOI:** 10.64898/2026.08.31.748276

**Authors:** Melis Gencel, Chloé Matta, Gisela Marrero Cofino, Sheela Ramanathan, Alfredo Menendez, Shimon Bershtein, Cang Hui, Adrian W.R. Serohijos

## Abstract

The outcome of ecological succession is often attributed to the characteristics of the invader or the resident community, but rarely to how the community’s interaction network reorganizes during assembly. Here, we track intraspecific lineage dynamics and infer time-resolved community interaction networks using Dynamic Covariance Mapping during ecological invasion of the mouse gut by a chromosomally barcoded, spectinomycin-resistant *Escherichia coli* K12 colonizer. The network is initially dominated by strong, predominantly inhibitory interactions, but as community diversity recovers, the distribution of interaction strengths contracts toward zero, producing a community increasingly dominated by weak and near-neutral interactions. The dominant eigenvalue of the DCM-inferred interaction matrix moves toward marginal stability predicted for dynamically assembling ecological networks. This pattern replicates across eight independent mice in two experimental cohorts, at both inter-and intra-species resolution. The ecological transition coincides with the reproducible resurgence of Paenibacillaceae to high relative abundance and persistent coexistence with *E. coli* under continued spectinomycin pressure. Whole-genome sequencing of recovered *Paenibacillus macerans* isolates identifies recurrent mutations in ribosomal protein S5 region associated with spectinomycin binding and strongly implicating this variant in resistance. Strikingly, under antibiotic pressure but without *E. coli* K12 invasion, resident Paenibacillaceae never blooms, indicating that expansion of the resistant population depends on the ecological context established by the colonizer. These findings show that gut microbiome succession is accompanied by a reproducible transition from strong toward weak interactions and link this network reorganization to the colonizer-dependent ecological benefit of antimicrobial resistance.

## Main text

Ecological succession, the ordered transformation of community composition following perturbation or colonization, has anchored ecological thinking since Clements^1^. Connell and Slatyer^2^ formalized this into three models (inhibition, tolerance, and facilitation). Subsequent succession theory has proposed that strong inhibitory interactions among competing pioneers can give way, as diversity accumulates, to weaker, more neutral or facilitative interactions^3,4^, a pattern tested in macroecological systems including intertidal^5^, plant^6^, and forest^7^ successions. By contrast, modern complex-network theory emphasizes interaction strength: stable, complex communities are promoted by networks of many weak interactions, with competitive structures often more stabilizing than strongly cooperative ones^8–12^. This is partly supported by microbiome diversity data in gnotobiotic mice, where resident competition determines whether an incoming strain establishes^13^, and *in vitro*, where interaction strength predicts invasion outcome in enteric^14^ and non-gut communities^15–17^. None of these studies, however, measured a whole-community interaction network reorganizing over an actual succession event in a living gut.

Which of these predictions holds in microbial communities, whether from classical succession or from complex-network theory, may depend less on diversity alone than on how the community arrived there. Assembly can depend critically on the order and timing of species arrival, a phenomenon termed priority effects^18,19^, generated by niche pre-emption or modification^18,20^ that broadly correspond to Connell and Slatyer’s inhibition and facilitation archetypes^2^. Modern coexistence theory formalizes when such effects could yield stable outcomes. Stabilizing mechanisms permit coexistence despite competition, while weak stabilizing mechanisms make outcomes increasingly dependent on fitness differences and arrival history^21^. Arrival order can itself determine whether a stabilizing niche difference is realized^22^. Gut establishment specifically depends on the niche overlap an incoming strain has with residents already present^23^. Whether a comparable dependence on colonization order and niche overlap governs the strength and sign structure of a reorganizing gut interaction network, rather than just strain-level establishment, remains untested.

Furthermore, priority effects and coexistence theory often treat the interactions themselves as fixed once an incoming strain establishes. In practice, ecological interactions are not fixed parameters but can evolve ^24–27^, adding an extra dimension to how a reorganizing network might change over time. Microbial systems are especially prone to rapid evolutionary change, given their short generation times, large population sizes, and exposure to strong, pervasive selection pressures such as resource competition, phage predation, and antimicrobials. Experimental evolution shows that competition alone can drive such change on ecologically relevant timescales^28,29^. Initially competitive pairs can evolve toward coexistence or facilitation through niche divergence or metabolic cross-feeding ^30–32^. Conversely, mutualistic or commensal interactions can be destabilized by evolution in fluctuating environments ^33^, and evolutionary change in one species can cascade through the community to alter interaction signs among multiple taxa ^34^.

The gut microbiome offers a tractable, ecologically structured system for testing these predictions. The gastrointestinal tract can be viewed as a habitat patch colonized through dispersal, selection, diversification, and drift ^35,36^. It transitions from an early, low-diversity, oxygen-tolerant pioneer community toward a stable, diverse anaerobic assemblage ^37–39^, and the stability of the mature community has been theoretically linked to networks of many weak, predominantly competitive interactions^40^. Prior interaction-network inference in the gut has generally fit a single, static network to an assembly trajectory,^41,42^ rather than testing whether network structure itself changes during succession. Most analyses have instead tracked compositional or functional change over time^38,39,43^ or used co-occurrence networks that cannot distinguish direct interactions from indirect, confounded associations^44^. Inferring time-resolved interaction structure from high-resolution temporal data offers a more quantitative route to testing which prediction holds^42,45,46^. Antimicrobial resistance evolution is a particularly important setting in which to examine eco-evolutionary dynamics: drug exposure alone can trigger emergent, community-level ecological behaviors not predictable from single-species responses^47^. Many resistance mutations have pleiotropic fitness effects whose sign and magnitude depend on environmental and competitive context ^48–50^. Their ecological effects cannot be inferred from fitness in isolation but instead depend on the identity and abundance of interacting community members^14,51–53^. Resistance evolution may therefore alter both the competitive position of the resistant lineage and the wider pattern of community interactions. Whether this depends on colonization order and niche overlap, as predicted above, has not been demonstrated.

Resolving these questions requires an approach that simultaneously tracks intraspecific lineage dynamics within a colonizing population and quantifies interspecific changes in community interactions over time. We combine chromosomal DNA barcoding^46,54–56^, which enables high-resolution tracking of hundreds of thousands of lineages within a colonizing bacterial population, with Dynamic Covariance Mapping (DCM). DCM is a dynamical systems framework for inferring signed, directional community interaction structure at both interspecific and intraspecific resolution^46^. Here, we apply this framework to characterize community succession in the mouse gut microbiome during invasion by a chromosomally barcoded, spectinomycin-resistant *E. coli* K12 colonizer, across eight independent mice. We find that the interaction network, initially dominated by strong and predominantly inhibitory interactions, progressively shifts toward a near-neutral interactions as community diversity recovers, while the dominant eigenvalue of the DCM-inferred interaction matrix moves toward the marginal-stability boundary, a pattern reproduced across all mice. This transition coincides with the reproducible expansion of resident Paenibacillaceae to high abundance and persistent coexistence with *E. coli* under continued spectinomycin pressure. Whole-genome sequencing of recovered *Paenibacillus macerans* isolates identifies a deletion in the ribosomal protein gene *rpsE* that is likely to perturb spectinomycin binding. Without *E. coli* K12 invasion, *Paenibacillaceae* remain at low abundance under antibiotic pressure alone, showing that expansion of the resistant population depends on the ecological context established by the colonizer. Together, these results suggest that ecological succession involves not simply species turnover but a reproducible reorganization from strong toward weak interactions, approaching marginal stability, while the ecological consequence of antimicrobial resistance depends on the organisms present within the assembling community.

## RESULTS

### High-resolution intra- and interspecific community dynamics during colonization under antibiotic selection

Because *E. coli* K12 colonizes poorly in the presence of a fully complex mouse gut microbiota, mice were pretreated for four weeks with a broad-spectrum antibiotic cocktail in drinking water (metronidazole 1 g/L, neomycin 1 g/L, ampicillin 1 g/L, and vancomycin 0.5 g/L), followed by a three-day antibiotic-free recovery period (Methods)^46^. Eight pretreated mice were then gavaged with a chromosomally barcoded *E. coli* K12 MG1655 library comprising approximately 10^8^ cells tagged with ∼10^6^ unique barcodes^46,55^. Each barcode comprised a 15-nucleotide degenerate sequence linked to the spectinomycin resistance gene *spR* (**Fig. 1a**), inserted at a neutral chromosomal locus (Tn7 site) and enabling individual lineages to be tracked by deep sequencing. Spectinomycin was maintained continuously in drinking water throughout the colonization experiment to maintain selection for the barcoded population. The mice were divided into two cohorts (cohort 1: m1–m4; cohort 2: m5–m8; **Fig. 1a**). The two cohorts were housed at separate facilities, and the experiments were conducted several months apart, providing an independent test of reproducibility across biological replicates. Fecal samples were collected at 3, 6, and 12 h post-gavage and daily thereafter for approximately two weeks. This enabled high-temporal-resolution reconstruction of *E. coli* lineage-frequency dynamics by deep sequencing of the chromosomal barcode locus (**Fig. 1c-d**; Methods). The same fecal samples were subjected to 16S rRNA gene sequencing to provide concurrent community-level profiling of the resident gut microbiota (**Fig. 1g-h**; Methods). *E. coli* abundance was additionally quantified by colony-forming unit plating on spectinomycin-selective agar (**Fig. 1b**).

**Figure 1:**
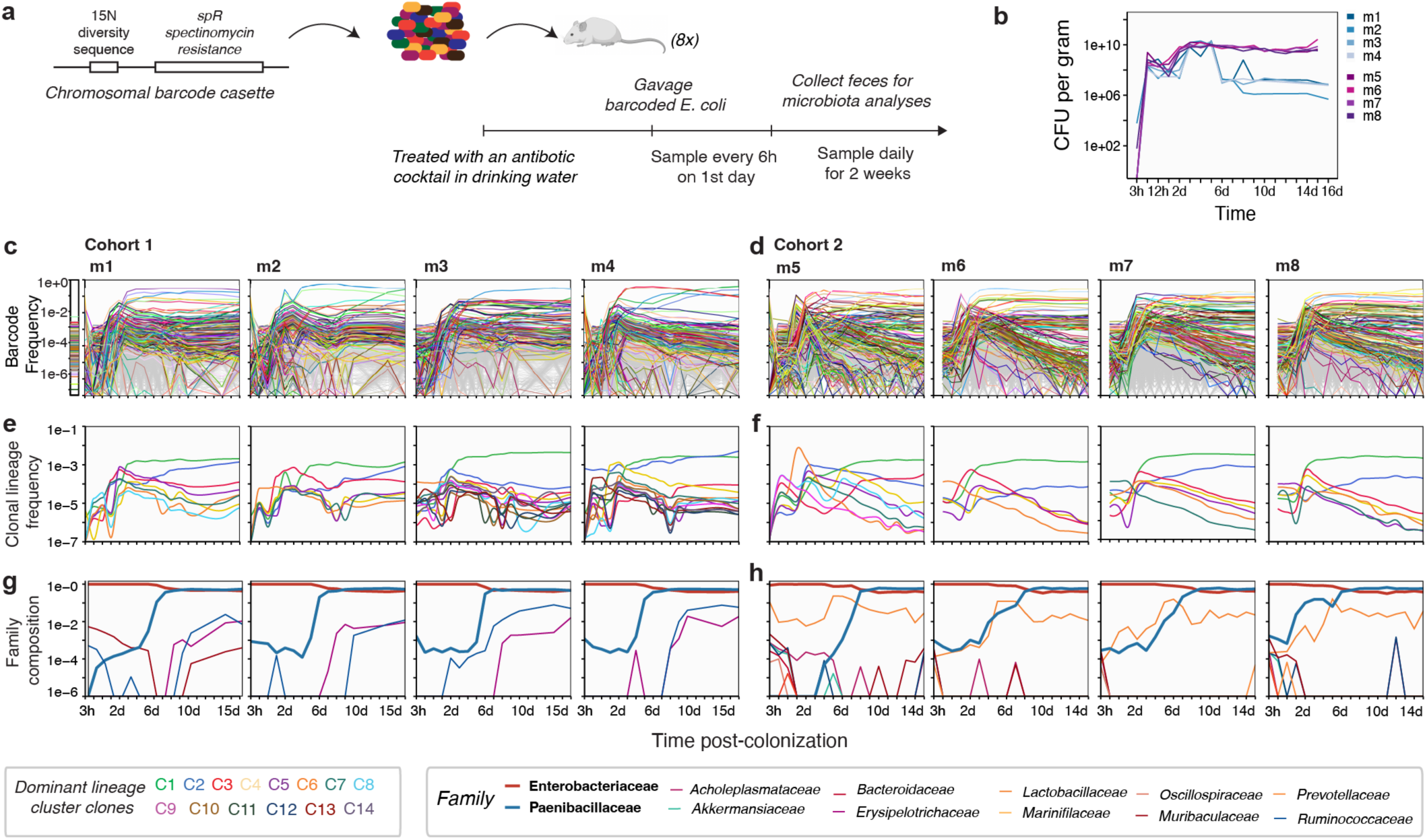
Colonization dynamics of spectinomycin-resistant and chromosomally barcoded *E. coli*. **(a)** Experimental design. The chromosomal barcode cassette comprises a 15-nucleotide degenerate barcode sequence flanked by a spectinomycin resistance determinant (*spR* gene), inserted at a neutral chromosomal locus (Tn7 site) and transmitted to all daughter cells, enabling lineage identification by deep sequencing. A library of ∼10⁸ barcoded *E. coli* K12 (MG1655) cells comprising ∼10⁶ barcodes was gavaged into eight antibiotic-pretreated mice (cohort 1: m1–m4; cohort 2: m5–m8). Fecal samples were collected at 3 h, 6 h, and 12 h post-gavage on day 1, then daily for two weeks, and subjected to barcode sequencing and 16S rRNA gene profiling. **(b)** Temporal dynamics of *E. coli* colonization, measured as colony-forming units (CFU) per gram of feces. **(c–d)** Barcode composition over time during colonization. The 1,000 most abundant barcodes are uniquely colored; all remaining barcodes are shown in gray. Color assignments are consistent across mice and cohorts to enable tracking of barcoded lineages. **(e–f)** Dominant *E. coli* lineage clusters in cohorts 1 and 2, identified using the Doblin^57^ clustering algorithm based on pairwise Pearson correlation of barcode frequency trajectories. These dominant clusters are ranked by their average frequency at the end of the experiment. **(g–h)** Microbial community composition profiled via 16S rRNA gene sequencing at the bacterial family level for cohorts 1 and 2.

*E. coli* successfully established in all eight mice, achieving high bacterial load during the initial days of colonization (**Fig. 1b**). Across both cohorts, the community dynamics exhibited two broadly reproducible phases: an initial phase of high *E. coli* abundance accompanied by a marked reduction in overall community diversity, and a subsequent succession phase marked by progressive recovery of microbial diversity and the expansion of Paenibacillaceae to high relative abundance (**Fig. 1g–h**). These phases were consistent in their ordering and qualitative features across all eight mice, despite the spatial and temporal separation of the two cohort experiments, indicating that the overall ecological transition was reproducible across independent experimental replicates.

Barcode frequency dynamics, visualized across the 1,000 most abundant lineages per mouse, revealed pronounced lineage turnover during the initial colonization phase, with distinct lineages rising and declining over the two-week observation window (**Fig. 1c–d**; **Extended Data Fig. 1a–b**). These dynamics were broadly similar between cohorts, indicating reproducible changes in lineage frequencies within the barcoded *E. coli* population. Barcode diversity computed using Hill diversity of order 1 (*^1^D*), equivalent to the exponential of Shannon entropy, declined sharply at early time points (**Fig. 2a**, purple line), consistent with a marked reduction in effective lineage diversity for the colonizing population (**Fig. 1c-d**). This early decline in barcode diversity coincided with the minimum in community diversity at the 16S level (**Fig. 2a**, orange line). As community diversity subsequently recovered, barcode diversity approached a plateau, coincident with the rise of Paenibacillaceae (**Fig. 2a**, blue line).

**Figure 2.**
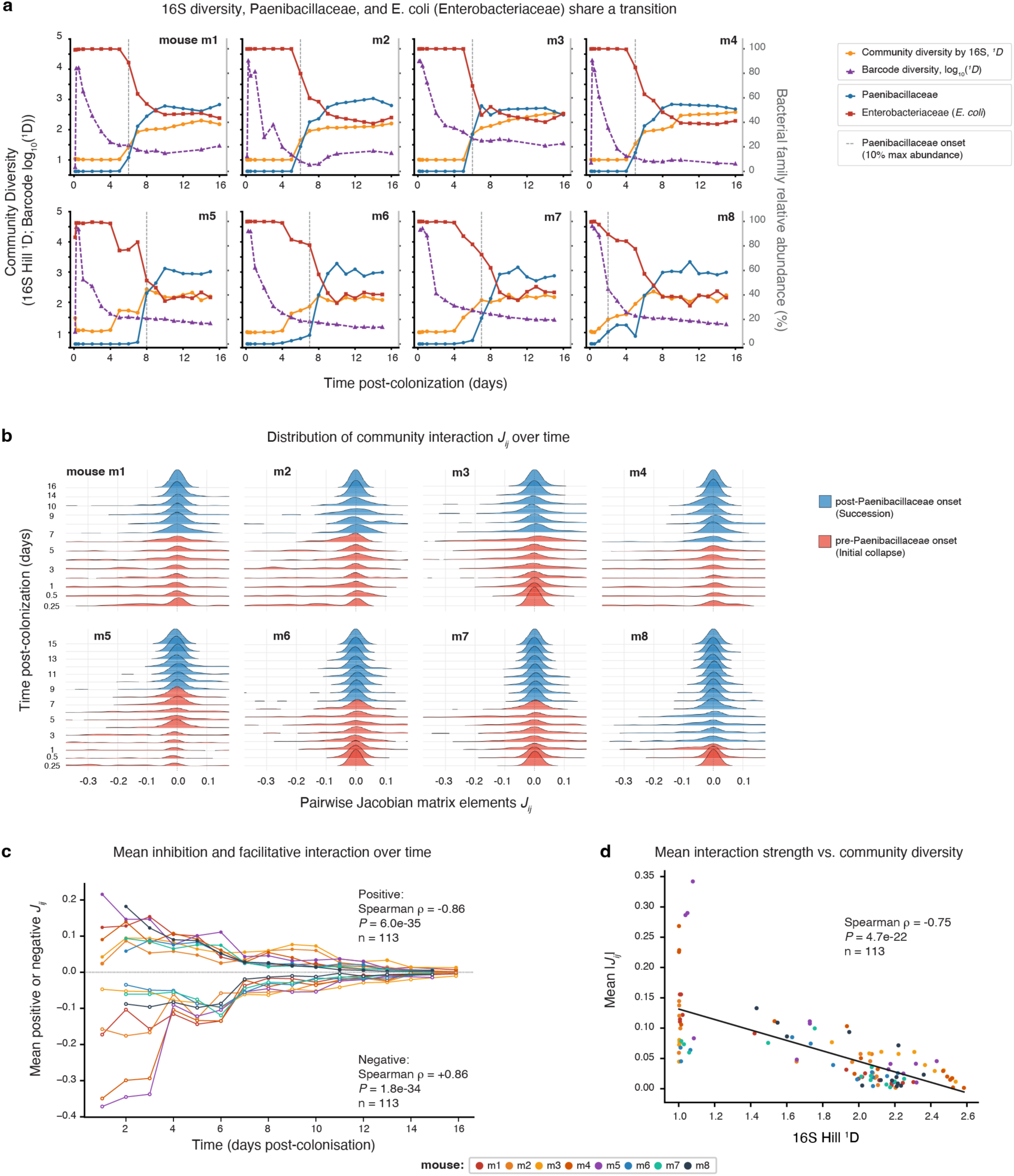
Ecological shift to near neutrality during succession tracks community diversity. **(a)** Time course of community diversity (16S Hill *¹D*, orange), *E. coli* barcode diversity (log_10_(*¹D*), purple), Paenibacillaceae relative abundance (blue), and Enterobacteriaceae (*E. coli*) relative abundance (red) for each of the 8 colonised mice (m1–m8). Vertical dashed gray line marks Paenibacillaceae onset (first timepoint ≥ 10% of per-mouse maximum Paenibacillaceae abundance). **(b)** Ridgeline distributions of pairwise DCM interaction elements *Jᵢⱼ* estimated from a 5-step sliding window, plotted on the real-day axis for each mouse. Red distributions (pre-Paenibacillaceae onset, initial collapse phase) are broad and left-skewed, reflecting a predominance of strong inhibitory interactions. Blue distributions (post-Paenibacillaceae onset, succession phase) progressively narrow and center near zero, indicating attenuation of initially strong inhibitory and facilitative coupling as the community assembles. **(c)** Mean positive (facilitative) and mean negative (inhibitory) *Jᵢⱼ* per mouse versus time post-colonisation. Both branches converge toward near-zero magnitude with time (positive: Spearman *ρ* = −0.86, *P* < 0.001; negative: *ρ* = +0.86, *P* < 0.001; *n* = 113 mouse-window observations, sliding window = 5). Colors denote individual mice (legend below panel). **(d)** Mean absolute interaction strength |*Jᵢⱼ*| versus community diversity (16S Hill ¹*D*), pooled across mice and sliding windows (*n* = 113). Interaction strength declines as diversity increases (Spearman *ρ* = −0.75, *P* < 0.001; black line, OLS fit). Points colored by mouse as in (c).

To identify recurrent patterns of intraspecific lineage dynamics from the high-dimensional barcode frequency data, we applied the Doblin clustering algorithm jointly to barcode trajectories from all eight mice^46,58^ (Methods). The number of lineage clusters was selected using the heuristic criterion implemented in Doblin, based on the relationship between distances among representative trajectories and the number of resulting clusters (**Extended Data Fig. 2a-b**). This procedure identified two dominant lineage clusters per mouse, designated C1 and C2, whose LOESS-smoothed representative trajectories captured the major temporal patterns of the *E. coli* population (**Extended Data Fig. 2c–d**; **Fig. 1e–f**). C1 and C2 each comprised a large and compositionally stable set of barcoded lineages across time, indicating that these clusters represented persistent lineage groupings rather than transient patterns generated by continual entry and loss of individual lineages.

Joint clustering of dominant *E. coli* lineage trajectories with bacterial family-level 16S abundance profiles revealed a consistent pattern across all eight mice: C1 and C2 clustered with Paenibacillaceae according to their temporal dynamics, while Enterobacteriaceae-associated families formed a distinct temporal profile (**Extended Data Fig. 3**). Bootstrap resampling with pvclust supported the stability of these groupings, with approximately unbiased (AU) values ≥95 for the group comprising Paenibacillaceae and the C1 and C2 lineage clusters across all animals (**Extended Data Fig. 3**) (Methods). This consistent joint clustering indicates that the temporal dynamics of the dominant *E. coli* lineage clusters were reproducibly associated with the rise of Paenibacillaceae during the community succession transition. This temporal coupling motivated subsequent analyses aimed at resolving the underlying interaction structure.

### Time-dependent interaction dynamics reveal a shift toward weak, near-neutral interactions during succession

To move beyond compositional tracking and quantify how the character of community interactions changed during colonization, we applied Dynamic Covariance Mapping (DCM)^8,46^ to infer time-dependent community interaction matrices from high-resolution abundance time-series for all eight mice. The DCM element *Jᵢⱼ* provides a covariance-based estimate of the directional interaction from taxon *j* to taxon *i*, linking the abundance of *j* to the rate of change (“growth rate”) of *i*, providing a signed, time-dependent measure of pairwise interaction strength that can be tracked as the community assembles^9,59^. Unlike static co-occurrence approaches, which identify associations but do not generally provide directed, time-resolved estimates of ecological interactions, DCM distinguishes inhibitory (negative) from facilitative (positive) inferred interactions and tracks their temporal dynamics, providing a framework to test predictions from classical succession^3–6^ and complex-network theories^8–11^ concerning changes in interaction sign and strength.

Two broad ecological phases were apparent across all mice, delineated by the timing of Paenibacillaceae onset (the first timepoint reaching 10% of each mouse’s peak Paenibacillaceae abundance), which ranged from day 2 to day 8 across mice (**Fig. 2a**, gray vertical line). During the initial collapse phase, before Paenibacillaceae onset, *E. coli* dominated the community, many resident bacteria declined to low relative abundance, and the distribution of pairwise *Jᵢⱼ* estimates computed from 5-step sliding windows was broad and left-skewed, reflecting a stronger contribution of inhibitory than facilitative interactions (**Fig. 2b**) (Methods). During the subsequent succession phase, as Paenibacillaceae expanded, the *Jᵢⱼ* distributions progressively narrowed and centred near zero, indicating systematic attenuation of both inhibitory and facilitative interactions as the community re-assembled (**Fig. 2b**). This transition in interaction structure was gradual and directional, and its timing coincided with the expansion of Paenibacillaceae observed in the compositional data. Consistent with this pattern, mean interaction magnitude declined systematically over the course of succession (**Fig. 2c**). The magnitudes of both positive (facilitative) and negative (inhibitory) *Jᵢⱼ* estimates declined toward zero with time post-colonization across mice (positive: Spearman *ρ* = −0.86, *P* < 0.001; negative: *ρ* = −0.86, *P* < 0.001; *n* = 113 mouse-window observations; **Fig. 2c**). Altogether, these results do not support a community-wide shift toward net facilitation^3–6^; instead, they reveal a progressive weakening of interactions toward near-neutrality, consistent with complex-networks predictions that community assembly is associated with increasingly weak interaction structure^8–11^.

Community diversity, derived from 16S rRNA gene profiles and quantified as Hill diversity of order 1 (*¹D*), showed a complementary temporal pattern across all eight animals (**Fig. 2a**). Diversity was low during the *E. coli* dominance phase, coinciding with the period of strongest inferred inhibitory interactions, and began recovering at its nadir, which preceded Paenibacillaceae onset in every mouse, by as little as one day to as much as six days. Interaction strength covaried with community diversity: mean absolute interaction strength |*Jᵢⱼ*|, pooled across mice and sliding windows, declined as diversity increased (Spearman *ρ* = −0.75, *P* < 0.001, *n* = 113; **Fig. 2d**).

The observed weakening of interaction strength was robust to the choice of temporal window used for the DCM estimation, as shown from the recomputed *Jᵢⱼ* across sliding windows spanning 2 to 9 consecutive sampling intervals (**Extended Data Fig. 4a**). The association between the shift toward near-neutral interactions and increasing community diversity was also robust to the choice of window size (mean absolute *Jᵢⱼ* vs. diversity, Spearman *ρ* = −0.58 to −0.75, *P* < 0.001; **Extended Data Fig. 4b**). The correlation was strongest at intermediate window sizes (5–6 steps, Spearman *ρ* = −0.75 and −0.74, respectively) and weakened at both extremes, consistent with different sources of uncertainty. At larger windows, the decline likely reflects fewer evaluable time points as the sliding window approaches the ends of each mouse’s sampling series (*n* = 129 at window 2 versus *n* = 81 at window 9), compounded by the window’s increasing tendency to average across the collapse-to-succession transition rather than resolve it. At smaller windows, the decline is consistent with greater estimation noise: with only 2–3 sampling intervals contributing to each local covariance estimate, *Jᵢⱼ* estimates showed substantially higher relative dispersion than at larger windows (coefficient of variation 2.8 at window 2 versus 1.5 at window 9), which may attenuate the apparent strength of the underlying relationship with diversity. The 5-step window used throughout the main text therefore provided a practical compromise between temporal resolution and estimation variability.

### Community approaches the marginal-stability boundary during succession

To determine whether the temporal shift in community interaction structure was accompanied by a change in local dynamical behavior, we tracked the dominant eigenvalue of the DCM-inferred community interaction matrix, Re(*λ*_max_), across sliding-window timepoints for all eight mice. For a system linearized around an equilibrium, local stability is determined by the real parts of the eigenvalues of the interaction matrix ^8–10,46^. The dominant value, Re(*λ*_max_), defines the stability boundary: negative values indicate asymptotic decay of small perturbations, whereas positive values indicate at least one locally divergent direction^8,12,60,61^. Because the communities sampled here were not assumed to be at equilibrium, we interpret this quantity more generally as an indicator of local dynamical convergence or divergence. Tracking this one dominant value at each timepoint, rather than every eigenvalue of every windowed matrix, provides a single matrix-level summary of these dynamics. It also avoids artificially inflating the sample size, since eigenvalues from the same matrix are not independent observations (Methods). Re(*λ*_max_) declined significantly with time post-colonization across all eight mice (**Fig. 3a**; Spearman *ρ* = −0.69, *P* < 0.001, *n* = 113 mouse-window observations, window = 5). It moved from strongly positive values early in colonization toward the marginal-stability boundary, Re(*λ*_max_) = 0, indicating progressively weaker local divergence. This pattern was robust to the choice of window size used to estimate the interaction matrices (**Extended Data Fig. 5**).

**Figure 3.**
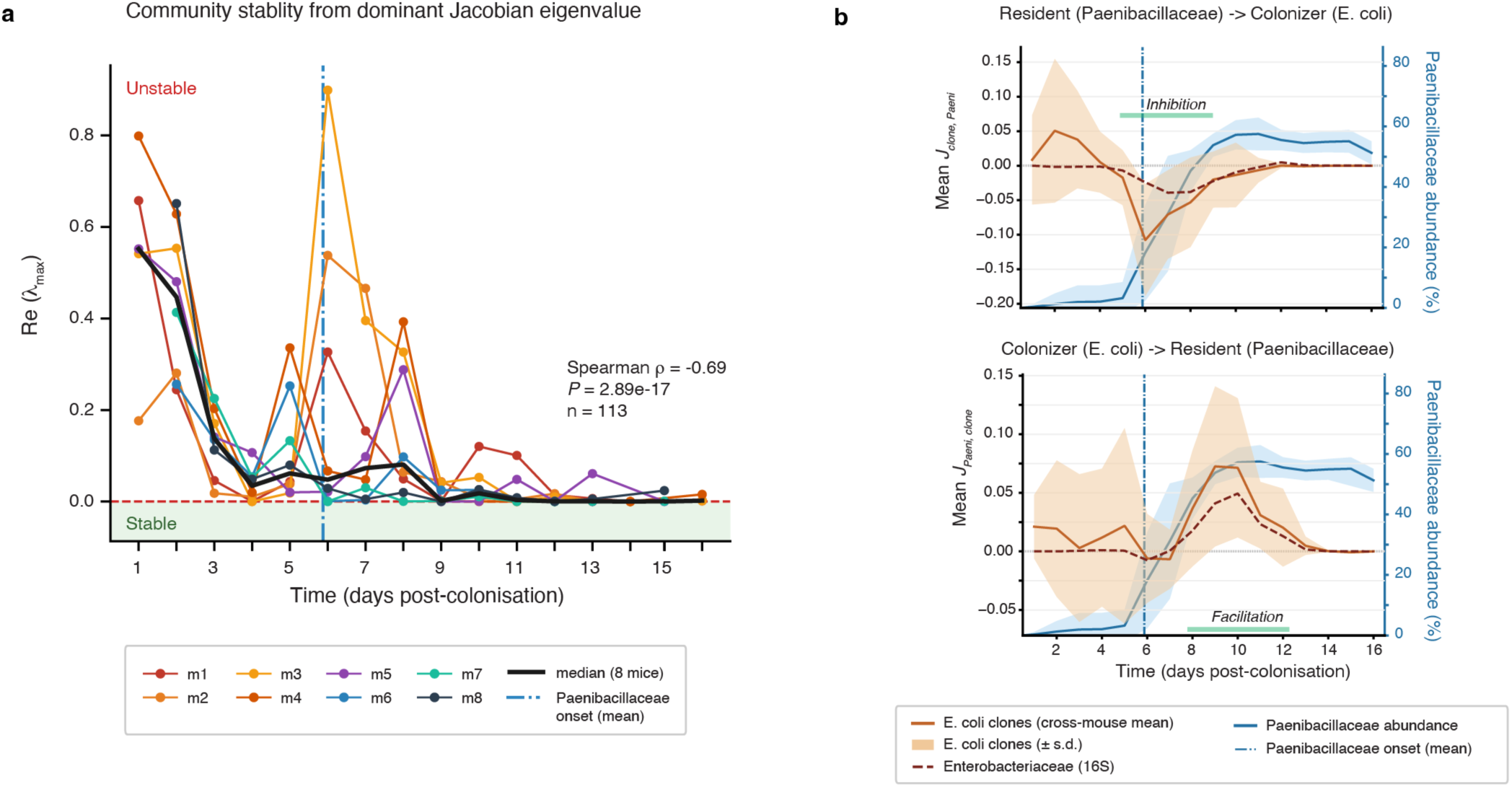
Community stability and colonizer–resident interaction dynamics during microbiome succession. **(a)** Dominant DCM interaction-matrix eigenvalue Re(*λ*_max_) (sliding window = 5) versus time post-colonization, one line per mouse (m1–m8, colors as in Fig. 2). Black line: cross-mouse median. Blue dash-dot line: mean Paenibacillaceae onset day (first day family abundance reaches 10% of its eventual maximum). Re(*λ*_max_) declines toward the marginal-stability boundary Re(*λ*_max_) = 0 (red dashed line), as succession proceeds (Spearman *ρ* = −0.69, *P* < 0.001, n = 113 mouse-window observations). **(b)** Directed pairwise interaction terms between the resident Paenibacillaceae and the *E. coli* colonizer population (chromosomally barcoded lineage clusters, aggregated), estimated by DCM; cross-mouse mean ± s.d. (shaded), left axis. Top: inferred interaction of Paenibacillaceae with the rat eof change of *E. coli* lineage clusters (mean *J_clone,Paeni_*) turns net negative around Paenibacillaceae onset (“Inhibition”, green bar), indicating the resident suppresses the colonizer as it establishes. Bottom: inferred interaction of *E. coli* lineage clusters with the rate of change of Paenibacillaceae (mean *J_Paeni,clone_*) turns positive following onset (“Facilitation”, green bar), consistent with the colonizer creating conditions to the resident’s rise. In both panels: solid blue line/right axis, Paenibacillaceae relative abundance (16S); dashed dark-red line/right axis, Enterobacteriaceae relative abundance (16S, shown for reference); blue dash-dot vertical line, mean Paenibacillaceae onset day (as in panel a). Together, (a) and (b) link the community-wide strong-to-weak interaction shift to a directed colonizer-resident feedback: an early positive inferred interaction from the colonizer to Paenibacillaceae is followed by increasingly negative interaction from Paenibacillaceae to the colonizer as the two populations move toward coexistence.

We interpret this trajectory as a shift toward marginal stability, a regime predicted for large, open ecological networks undergoing persistent assembly and turnover. In such systems, continued species turnover can maintain the leading eigenvalue near the boundary between locally convergent and divergent dynamics rather than driving the network deep into the stable regime^8,12,60,61^. The declining Re(*λ*_max_) trajectory is consistent with the community moving toward this marginal regime as succession proceeds, mirroring the shift toward weaker, near-neutral interactions described above (**Fig. 2**).

This overall decline was punctuated by two distinct episodes of markedly elevated Re(*λ*_max_), indicating transient increases in local divergence rather than a smooth monotonic approach toward the boundary. The first and largest coincided with the earliest days of colonization, when *E. coli* invasion under antibiotic pressure coincided with the sharpest community collapse (median Re(*λ_max_*) reaching 0.55 within the first two days across mice). A second, smaller resurgence occurred around days 5-6, coincident with Paenibacillaceae onset, and was most pronounced in cohort 1 (m1-m4, particularly m2 and m3; **Fig. 3a**). This is consistent with a second, transient increase in local divergence as Paenibacillaceae expands and the interaction network reorganizes, followed by a return toward the low-eigenvalue regime. This pattern (an early collapse-associated peak, a second Paenibacillaceae-associated peak, and an overall decline toward the marginal-stability boundary) was reproducible across cohort 1 (m1-m4) and cohort 2 (m5-m8), which were colonized and profiled in independent experiments conducted months apart. This recurrence indicates that the pattern was reproducible across independent experimental cohorts rather than being specific to a single experimental run.

Notably, Re(*λ*_max_) approached but did not cross zero within the two-week observation window. By day ≥13, the median across all mice was 0.0008, essentially at the boundary. Every late-window observation remained marginally positive, and none crossed into the negative regime. Thus, the communities approached the marginal-stability boundary closely but did not enter the formally stable, negative eigenvalue regime during the observation period. The community had been substantially perturbed by antibiotic pretreatment and *E. coli* invasion, and the interaction-strength distribution identified above was itself still narrowing at the end of the observation window (**Fig. 2**). These trajectories are consistent with continued relaxation toward a marginal regime, although longer observation would be required to determine whether the community ultimately crosses the boundary, stabilizes near it, or continues to fluctuate around it.

### Directional colonizer-resident interaction dynamics

To characterize the directional structure underlying the broader strong-to-weak interaction transition at the level of specific colonizer–resident pairs, and to resolve the relative contributions of each partner to the interaction dynamics, we computed the interspecific DCM interaction elements linking *E. coli* lineage-cluster trajectories with Paenibacillaceae abundance across all eight mice. Specifically, the quantity *J*_cluster,Paeni_, which estimates the directed interaction from Paenibacillaceae to the rate of change of *E. coli* lineage clusters, became negative from approximately day 5 onward, reaching its most negative mean value at day 6 (mean = −0.11, averaged across *E. coli* lineage clusters and mice), and returning towards zero by approximately day 12 (**Fig. 3b**, top panel). This temporal profile is consistent with an inhibitory interaction of the expanding Paenibacillaceae population with *E. coli* during the early succession phase. The corresponding DCM interaction element involving family-level Enterobacteriaceae trajectory derived from 16S rRNA gene profiling showed a qualitatively similar trajectory (**Fig. 3b**, top panel, maroon dashed line), providing a concordant community-level signal for the negative association observed at *E. coli* lineage resolution.

Conversely, *J*_Paeni,cluster_, which estimates the directed interaction from *E. coli* lineage clusters to the rate of change of Paenibacillaceae, became positive during the same temporal window (**Fig. 3b**, bottom panel), indicating a positive inferred interaction of *E. coli* with *Paenibacillaceae* expansion. The positive *J*_Paeni,cluster_ signal preceded the strongest negative *J*_cluster,Paeni_ values, revealing a temporally ordered asymmetry in the inferred interaction between the two populations. This sequence is consistent with a scenario in which *E. coli* initially creates or modifies conditions that favor Paenibacillaceae expansion, either directly through changes in the local nutritional or chemical environment or indirectly through effects on other resident taxa. The subsequent increase in Paenibacillaceae abundance is, in turn, associated with an increasingly interaction on the *E. coli* population. These reciprocal DCM estimates are therefore consistent with an asymmetric ecological feedback, whereby an early positive interaction of the colonizer with Paenibacillaceae is followed by an increasingly negative interaction of *Paenibacillaceae* with the colonizer as succession proceeds.

Resolving the *E. coli* population at lineage level also revealed interaction dynamics of greater magnitude than those apparent from family-level 16S rRNA gene profiles. The *Paenibacillaceae*–*E. coli* lineage-level interaction estimates were consistently larger in absolute magnitude than the corresponding *Paenibacillaceae*–Enterobacteriaceae estimates derived from 16S data (**Fig. 3b**). Thus, lineage-resolved barcoding captured stronger temporal interaction signals than were apparent after aggregation at the bacterial family level.

### Colonizer dependence of the community interaction shift during succession

To determine whether the shift toward weak, near-neutral interactions observed in colonized mice depended on *E. coli* colonization, rather than being a generic consequence of antibiotic exposure in the resident microbiota alone, we analysed a control cohort of four mice (cm1-cm4). These mice received the identical four-week antibiotic cocktail followed by the same three-day recovery period and subsequent spectinomycin treatment, but were not gavaged with barcoded *E. coli* (**Fig. 4**). This control group provided a direct ecological comparison. If antibiotic treatment of the resident microbiota alone were sufficient to generate the succession dynamics observed in colonized mice, similar community trajectories and Paenibacillaceae abundance would be expected in the presence and absence of the *E. coli* colonizer.

**Figure 4.**
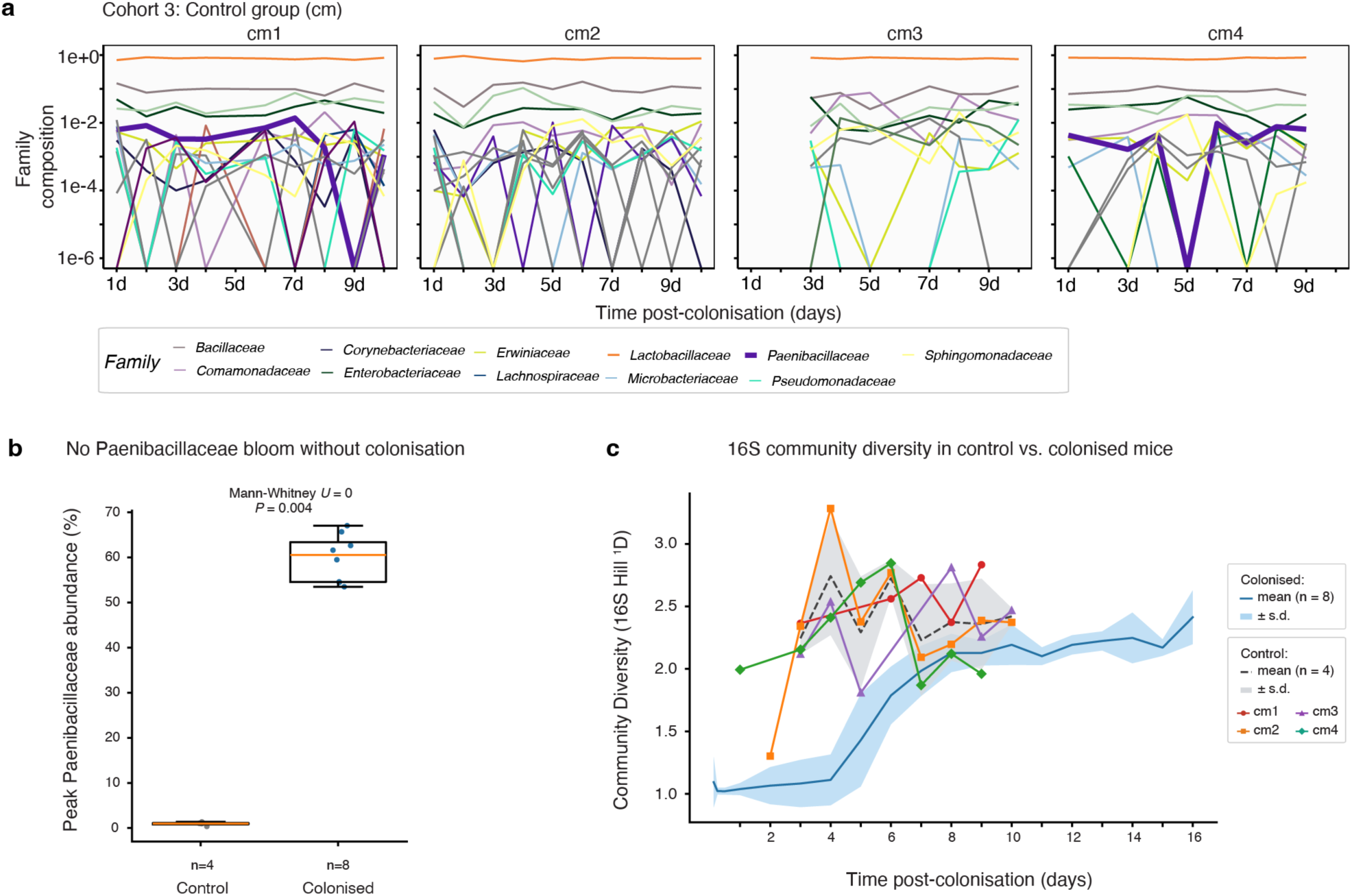
Antibiotic-treated control mice do not recapitulate the community collapse or bloom of *Paenibacillaceae* in colonised mice. (a) 16S family-level relative abundance (log scale) over the first ∼10 days, one panel per control mouse (cm1–cm4, “Cohort 3”). Paenibacillaceae (bold purple) is highlighted against all other detected families; it remains a minor, fluctuating community member throughout (frequently dropping to the detection floor) while Lactobacillaceae dominates near saturation in every mouse. **(b)** Peak Paenibacillaceae relative abundance per mouse, control versus colonized (one value per biological replicate, not per timepoint). Peak abundance is completely separated between groups: every control mouse tops out under 1.4%, every colonized mouse peaks above 53% (Mann-Whitney *U* = 0, *P* = 0.004; *n* = 4 control, *n* = 8 colonized), showing that, under the experimental conditions tested, Paenibacillaceae does not bloom without *E. coli* colonization. **(c)** 16S community diversity (Hill ¹*D*) versus time post-colonization, control mice (grey, dashed mean ± s.d., individual cm1–cm4 traces) versus colonized mice (blue, solid mean ± s.d., *n* = 8). Diversity in colonized mice collapses immediately post-colonization and only recovers toward control-range values by ∼day 14–16, whereas control-mouse diversity stays elevated throughout, consistent with the diversity loss depending on *E. coli* colonization rather than antibiotic exposure alone.

The control mice exhibited a markedly different ecological trajectory from the colonized animals. All four control animals showed relatively stable community diversity throughout the ten-day observation window (16S-derived Hill diversity *¹D* ranging from 1.3 to 3.3 across mice and timepoints), with Lactobacillaceae remaining the dominant family throughout the experiment and no acute collapse in diversity at any timepoint (**Fig. 4a**). The difference between groups was most direct when comparing the peak *Paenibacillaceae* abundance achieved by each animal, one value per biological replicate rather than per timepoint. Across the ten-day observation window, no control mouse exceeded 1.4% relative Paenibacillaceae abundance (range 0.4-1.4%, n = 4), while every colonized mouse exceeded 53% (range 53.5-67.0%, *n* = 8), with complete separation between groups (Mann-Whitney U = 0, *P* = 0.004; **Fig. 4b**). This contrast indicates that, under the experimental conditions used here, *Paenibacillaceae* expansion was contingent on *E. coli* colonization rather than on antibiotic exposure alone.

The overlay of control diversity trajectories on the colonized mean ± s.d. further illustrated this difference. Even at their lowest, control mice did not approach the low community diversity (*¹D* ≈ 1) characteristic of the initial collapse phase in the eight colonized animals (**Fig. 4c**). These results demonstrate that antibiotic exposure in the absence of the *E. coli* colonizer did not reproduce the diversity collapse or Paenibacillaceae expansion that characterized the successional transition in colonized mice. *E. coli* colonization therefore emerges as a necessary component of this succession sequence under the experimental conditions tested.

### Convergent lineage and community dynamics across mice suggest reproducible ecological selection

The ecological shift we describe was reproducible across all eight mice, spanning two cohorts, two facilities, and experiments conducted months apart, raising the question of whether this convergence reflects selection on a shared underlying genetic mechanism. To test this, we jointly clustered dominant *E. coli* lineage trajectories and bacterial family-abundance profiles using shape-based distance (SBD), which compares the shape of time series independent of absolute amplitude. This resolved three broad trajectory groups present in both cohorts (**Extended Data Fig. 6a-b**), with pairwise SBD values consistently lower within than between groups (*P* < 0.001; **Extended Data Fig. 6d-e**). The most notable group linked Paenibacillaceae with the two dominant *E. coli* lineage clusters (C1 and C2), consistent with the temporal coupling already evident from the *Jᵢⱼ* analyses, and this grouping was supported by bootstrap resampling (AU ≥ 95 across all eight mice; **Extended Data Fig. 3**). Beyond trajectory shape, dominant lineage clusters within this group also shared significantly more barcode identity across mice within the same cohort than expected by chance (Szymkiewicz-Simpson overlap; **Extended Data Fig. 6c,f**), indicating that some of the same barcoded *E. coli* lineages were repeatedly selected from the ∼10⁶-barcode inoculum across independent animals. Together, this convergence, lineage-level dynamics, and community trajectories, and barcode identity across eight independent mice points to reproducible ecological selection, raising the possibility of a shared genetic mechanism, and motivated a direct genomic investigation of succession-phase Paenibacillaceae isolates.

### Whole-genome sequencing identifies an *rpsE* deletion associated with spectinomycin resistance in gut-derived *Paenibacillus macerans*

Paenibacillaceae resurgence under continuous spectinomycin pressure indicated that the succession-phase population could tolerate spectinomycin exposure. To characterize this population, we sought to isolate Paenibacillaceae from fecal samples of cohort 2 mice (m5– m8) by heat treatment, exploiting the endospore-forming capacity of *Paenibacillus* to enrich for heat-resistant cells while reducing non-sporulating microbes, including the barcoded *E. coli* (Methods). Heat-treated day-12 samples were plated on LB agar containing spectinomycin (100 μg/ml), whereas samples from earlier time points (3–12 h) were plated on antibiotic-free LB agar, followed by anaerobic incubation for 48 h. Visible colonies were recovered from mice m5, m6, m7, and m8 in the day-12 samples; however, no colonies were recovered from the earlier time points. Sanger sequencing of the 16S rRNA gene identified the recovered colonies as *Paenibacillus*, with subsequent whole-genome sequencing assigning the isolates to *P*. *macerans*. The isolates displayed a characteristic filamentous colony morphology across the tested LB agar concentrations (**Figure 5a**).

**Figure 5.**
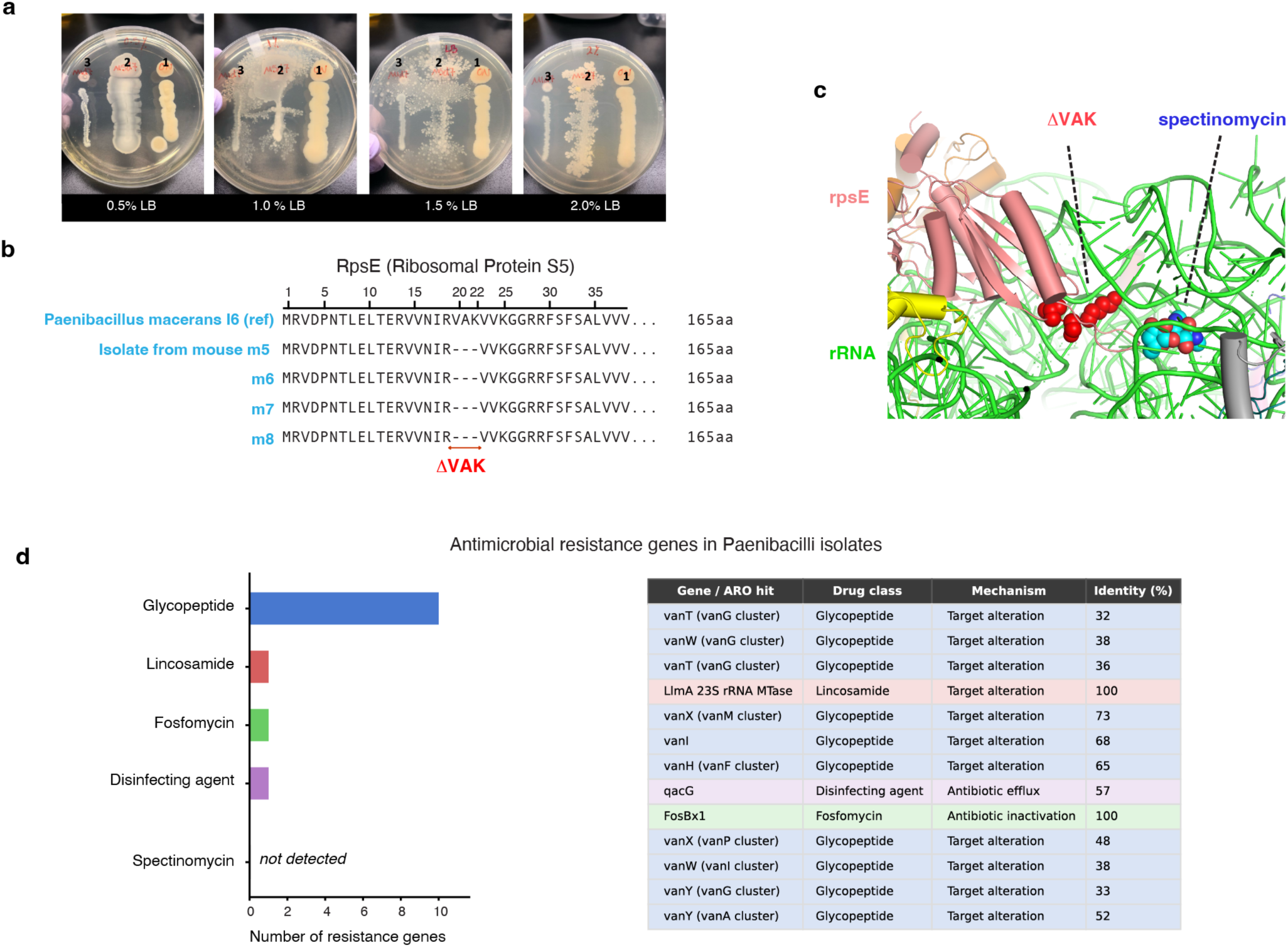
Whole-genome sequencing identifies a chromosomal *rpsE* deletion conferring spectinomycin resistance in gut-adapted *Paenibacillus macerans* isolates. (a) Colony morphology of a *P. macerans* gut isolate grown on LB agar at varying nutrient concentrations (0.5–2.0% LB), displaying the characteristic filamentous morphology of this species. **(b)** Protein sequence alignment of RpsE (30S ribosomal protein S5) from the *P. macerans* I6 reference strain and isolates recovered from the gut of colonized mice (m5– m8). All gut isolates carry an in-frame deletion of three consecutive residues (ΔVAK, positions 20–22) within the N-terminal β-hairpin. **(c)** Structural context of the ΔVAK deletion mapped onto the E. coli 70S ribosome (Protein Databank ID: 4V56). The deleted residues (red) lie at the interface between ribosomal protein S5 (rpsE, pink) and 16S rRNA helix h34 (green), within the spectinomycin-binding pocket (cyan). Loss of this contact is predicted to disrupt antibiotic binding without impairing core ribosomal function. **(d)** Antimicrobial resistance gene survey of the *P. macerans* I6 genome against the CARD database (RGI, Strict threshold). Detected genes fall within glycopeptide, lincosamide, fosfomycin, and disinfecting-agent classes; no spectinomycin resistance determinants were identified, confirming that resistance in gut isolates is attributable solely to the chromosomal ΔVAK mutation.

Because the colonizing *E. coli* harbored the spectinomycin-resistance cassette *spR* within its chromosomal barcode construct, we first tested whether the resistance phenotype in *P. macerans* could reflect acquisition of this cassette from the colonizer (**Fig. 1b**). This possibility was motivated by the sustained coexistence of Paenibacillaceae and *E. coli* at high relative abundance in the experiment (**Fig. 1g-h**; **Fig. 2a**). Thus, we performed whole-genome sequencing (WGS) and comparative genomic analysis against the *P. macerans* I6 reference genome (Methods). However, we did not find the *spR* cassette in all available *P. macerans* isolates, ruling out horizontal acquisition of the colonizer-derived *spR* cassette in these isolates and prompting investigation of chromosomal resistance-associated variation. Among the variants shared by the four gut-derived isolates, we identified a nine-nucleotide in-frame deletion in *rpsE*, encoding 30S ribosomal protein S5, that removed three consecutive residues (valine V20, alanine A21, and lysine K22) from the

N-terminal β-hairpin (ΔVAK; **Fig. 5b**). The deletion was identical in sequence and position across all four isolates recovered from mice m5-m7, indicating reproducible enrichment of the same resistance-associated genotypes across these isolates. The location of ΔVAK provided a structural basis for spectinomycin resistance. We mapped the affected residues onto the *E. coli* 70S ribosome co-crystallized with spectinomycin (PDB 4V56; **Fig. 5c**). The N-terminal β-hairpin of *RpsE* (residues 20–22) contacts helix h34 of 16S rRNA, which contains the primary spectinomycin-binding site, with K22 making direct electrostatic contact with the phosphate backbone within the drug-binding pocket. Deletion of the VAK triplet disrupts both this contact and the β-hairpin geometry that positions K22, thereby perturbing the structural scaffold that spectinomycin exploits for binding. Notably, this disruption occurs in a peripheral loop rather than the ribosomal core, consistent with the robust growth of gut isolates across nutrient conditions (**Fig. 5a**) and previous reports of limited fitness costs associated with *rpsE* resistance mutations under non-selective conditions^49,50,62,63^.

In addition, comprehensive screening of the *P. macerans* isolate genomes against the CARD (Comprehensive Antibiotic Resistance Database)^64^ identified homologs associated with resistance of four antibiotic classes (glycopeptide, lincosamide, fosfomycin, and disinfectants/biocides) with sequence identities ranging from 32% to 100% (**Fig. 5d**), consistent with intrinsic chromosomal homologs. No known spectinomycin-resistance determinant was detected by CARD analysis. Together with the location of ΔVAK in a region of S5 previously implicated in spectinomycin resistance, this result supports the *rpsE* deletion as the leading candidate genetic basis of the resistance phenotype.

The repeated recovery of the identical ΔVAK-bearing *P. macerans* genotype under sustained spectinomycin exposure is consistent with positive selection for this variant, although the present data do not distinguish *de novo* mutation from selection on pre-existing variation. Ribosomal resistance mutations can carry context-dependent fitness effects whose sign and magnitude depend critically on the competitive environment^49,50^. Consistent with such context dependence, Paenibacillaceae remained absent or at trace abundance in antibiotic-treated mice not colonized by *E. coli* (**Fig. 4**), whereas it expanded to high relative abundance in all eight *E. coli*-colonized animals. Thus, expansion of the resistant population was contingent on the ecological context established by the *E. coli* colonizer, potentially through competitive release, nutritional modification, or alteration of the gut antibiotic gradient, linking chromosomal resistance evolution to the colonizer-dependent eco-evolutionary dynamics of the gut succession transition.

## DISCUSSION

### Demonstrating a shift toward near-neutral, not facilitative, interaction structure

The strong-to-near-neutral shift we document does not support a community-wide drift toward facilitation, as emphasized in classical succession framework and subsequently developed through facilitation theory ^1–4^, which predict that the balance of ecological interactions should shift toward increasingly cooperative, positive interactions as early colonizers are replaced by later-successional taxa. Instead, both the positive and negative branches of the pairwise interaction distribution weakened toward zero (**Fig. 2c**), and mean absolute interaction strength declined as community diversity recovered (**Fig. 2d**), a pattern that supports the alternative prediction from complex-networks theory that persistent, diverse communities are organized around weak, near-neutral interactions rather than a drift toward net facilitation ^8–11^. This interpretation is consistent with recent demonstrations that resident competition governs strain establishment in gnotobiotic mice ^13^ and that interaction strength and arrival order jointly determine invasion outcome in vitro ^15,16^. This highlights, in the gut microbiome specifically, the contrast between these predictions for how succession reshapes interaction structure. We also demonstrate the shift at the level of individual pairwise community interaction estimates, revealing a structured, temporally ordered reorganization of the community interaction network, rather than a simple change in the overall magnitude of interactions. Prior work has largely characterized gut microbiome interactions from compositional changes or co-occurrence associations, which do not generally resolve interaction sign, direction, or dynamics ^38,39,43^; time-series matrix inference has been attempted ^42,45^ but without simultaneously resolving intraspecific dynamics or evidence of a directional successional shift in interaction structure. By integrating intraspecific barcoding with community interaction inference across eight independent replicates, our approach resolves both dimensions simultaneously, establishing the interaction shift as a reproducible phenomenon across individual animals rather than a statistical tendency from pooled community-level analyses.

### A colonizer-mediated route to Paenibacillaceae resurgence

The sequential structure of the colonizer-resident DCM interaction profiles, in which an early positive inferred interaction of *E. coli* with Paenibacillaceae is followed by an increasingly negative interaction of Paenibacillaceae with the colonizer, is a directed, pairwise finding distinct from the network-wide shift toward near-neutral interaction structure described above. This directed relationship instead suggests a specific, colonizer-mediated route by which *E. coli*’s presence contributes to the succession transition. One plausible route for this apparent competitive release is *E. coli*-mediated suppression of Lactobacillaceae. In control mice not receiving the colonizer, Lactobacillaceae remained the dominant taxon across all animals, whereas Paenibacillaceae remained at very relative abundance or was undetected (**Fig. 4a**); in colonized cohort 2 mice, *Lactobacillaceae* increased only after Paenibacillaceae had established and never reached comparable abundance (**Fig. 1h**). This pattern is consistent with *E. coli*-mediated suppression of Lactobacillaceae, potentially relieving competitive constraints on Paenibacillaceae and enabling its subsequent resurgence. Consistent with this interpretation, the absence of pronounced Paenibacillaceae expansion and diversity collapse in all four control mice indicates that *E. coli* colonization is a necessary component of this ecological transition under the experimental conditions (**Fig. 4**). This has direct implications for colonization resistance: the resident microbiota’s ability to re-establish community structure following perturbation may depend not only on its intrinsic recovery trajectory but also on specific ecological interactions with the colonizer that create or modify the conditions for the resurgence of resident taxa.

### Antimicrobial resistance as an eco-evolutionary driver of succession

The context dependence of *Paenibacillaceae* expansion, together with the identification of the *rpsE* ΔVAK variant in resistant *P. macerans* isolates from cohort 2, illustrates a principle with implications beyond gut ecology. Resistance mutations are typically framed as adaptations to antimicrobial pressure, but their ecological consequences depend critically on biotic context^49,50^. Here, the resistant *P. macerans* population persisted under spectinomycin selection, whereas its expansion to high abundance occurred only in the ecological context established by the *E. coli* colonizer, potentially through competitive release or other forms of niche modification. This parallels recent evidence that antibiotic exposure itself can trigger emergent, community-level ecological behaviors not predictable from single-species responses^47^, reinforcing that both the perturbation and the resulting resistance phenotypes act on the community as a whole rather than on isolated lineages. This eco-evolutionary coupling positions antimicrobial resistance as a potential driver of community-level ecological dynamics^14,48,53^. The ΔVAK deletion may therefore contribute, through its putative role in spectinomycin resistance and the resulting expansion of *P. macerans*, to the succession transition and associated restructuring of the community interaction network. The ecological consequences of antibiotic use may therefore extend beyond direct effects on susceptible taxa to include resistance-driven succession dynamics whose sign and magnitude are contingent on the biotic environment.

### Limitations and future directions

Several limitations qualify these conclusions. First, DCM infers local interaction structure using a first-order linear approximation and requires interactions to remain sufficiently stable within each estimation window ^8,46^. During the rapidly changing early colonization phase, *Jᵢⱼ* values are therefore best interpreted as inferred interaction estimates rather than direct measurements of causal effects ^65,66^. DCM also captures pairwise covariance-based interaction structure, so it may under-detect facilitation operating through higher-order or indirect pathways such as cross-feeding. The near-neutral shift we observe describes this dominant, pairwise inferred signal, and does not rule out subtler facilitative mechanisms operating alongside it. Second, our stability inference relies on the dominant eigenvalue Re(*λ*_max_) at each sliding window, rather than the full spectrum, both to avoid pseudoreplication and because Re(*λ*_max_) the spectral abscissa is the relevant eigenvalue summary for local asymptotic stability of a linearized equilibrium. This estimate depends on the fidelity and dimensionality of matrices resolving only 9-17 taxa per mouse, substantially fewer than an intact gut microbiome. The sampled communities were also not necessarily at equilibrium, so changes in Re(*λ*_max_) are best read as shifts in local dynamical convergence or divergence rather than formal proof of stability. Consistent with this, Re(*λ*_max_) approached but did not cross the marginal-stability boundary within our two-week window. Whether it eventually does, or instead remains close to that boundary, as predicted for some persistent open networks, remains untested and would require longer-term sampling. Third, the mice were also deliberately pretreated with a broad-spectrum antibiotic cocktail to permit *E. coli* colonization. This simplification alters the ecological context and may limit generalization to intact communities. Future work using gut-adapted or pathogenic strains capable of colonizing microbiotas of greater baseline complexity could test whether the same transition holds under more physiologically intact conditions. Fourth, whole-genome sequencing was restricted to cohort 2 isolates. Whether the identical ΔVAK deletion was also present in resistant cohort 1 Paenibacillaceae could not be confirmed. Genomic and structural evidence strongly implicates ΔVAK in spectinomycin resistance, but direct functional validation remains a priority for future work.

## METHODS

### Chromosomal barcoded E. coli

Chromosomally barcoded *E. coli* populations were generated using the Tn7 transposon system as previously described^55^. Briefly, integration plasmids carrying the barcode and spectinomycin resistance cassettes were purified from TransforMax EC100D *pir*+ cells (Lucigen) and introduced into *E. coli* K12 MG1655 pre-loaded with the Tn7 helper plasmid, with transposase expression induced by arabinose. Transformants were selected on LB agar with 100 μg/ml spectinomycin, verified for chromosomal integration by PCR at the Tn7 insertion site, then scraped, pooled, and stored at −80°C in 15% glycerol. The resulting library, comprising approximately 10^6^ barcodes, was administered to mice by oral gavage at approximately 10^8^ cells per animal (**Fig. 1a**).

### Mouse gut colonization experiment

Colonization experiments were conducted at two independent facilities at Université de Sherbrooke following identical protocols, with four mice per facility (cohort 1: m1–m4; cohort 2: m5–m8; cohorts run months apart). To reduce the resident gut microbiota, mice received a broad-spectrum antibiotic cocktail in their drinking water consisting of metronidazole (1 g/L), neomycin (1 g/L), ampicillin (1 g/L), and vancomycin (0.5 g/L) for four weeks (**Fig. 1a**). Following a three-day antibiotic-free recovery period, mice were orally gavaged with approximately 10^8^ cells of the chromosomally barcoded *E. coli* MG1655 library (∼10^6^ barcodes) as described above. A control group (cm1–cm4) received the same antibiotic regimen but was not gavaged with *E. coli* to assess the effect of antibiotics alone on community composition (**Fig. 4**). Fecal samples were collected at 3, 6, and 12 hours post-gavage, then daily through day 16 for cohort 1, day 15 for cohort 2, and day 10 for the control group. Samples were plated on LB agar with spectinomycin (50 μg/ml) to quantify *E. coli* abundance; additional aliquots were stored at −80°C for microbiota analysis.

Genomic DNA (gDNA) was extracted from whole fecal pellets using the QIAamp Fast DNA Stool Mini Kit (Qiagen; cat. no. 51604) and served as the template for two downstream workflows: chromosomal barcode amplification and 16S rRNA gene sequencing.

### Barcode amplification

To amplify chromosomal barcodes and incorporate Illumina adapter sequences, we employed a two-step PCR strategy using PrimeSTAR GXL DNA Polymerase (Takara, Cat: R050B). For the first PCR, 200 ng of gDNA per sample was used as the template. Thermal cycling consisted of an initial denaturation at 95°C for 5 min; 40 cycles of denaturation at 95°C for 10 s, annealing at 55°C for 15 s, and extension at 68°C for 45 s; and a final extension at 68°C for 5 min, followed by a hold at 4°C. PCR products were purified using the NucleoSpin Gel and PCR Clean-up Kit (Takara) to remove residual primers and reaction components. For the second PCR, Nextera XT primers Set A (96 Indexes, 384 Samples; cat. no. FC-131-2001) and PrimeSTAR GXL DNA Polymerase were used to add sample-specific Nextera indices. Thermal cycling consisted of an initial denaturation at 94°C for 5 min; 12 cycles of denaturation at 95°C for 10 s, annealing at 55°C for 15 s, and extension at 68°C for 45 s; and a final extension at 68°C for 5 min, followed by a hold at 4°C. The final indexed PCR products were purified using magnetic beads (Beckman Coulter) to remove excess reaction components. The purified libraries were pooled and supplemented with 15% PhiX DNA to improve sequence complexity.

### Barcode extraction from raw sequencing data

To extract barcode sequences from raw sequencing data, we first filtered reads by quality, discarding reads with an average Phred score below 30. Barcodes were identified and extracted using BarcodeCounter2^67^, which leverages BLASTn+ v2.6.0 ^68,69^ to detect barcode sequences based on a predefined template. Due to errors introduced during barcode synthesis, sequencing, and library preparation, extracted barcode lengths varied. Although the barcode design comprised 15 random nucleotides (15N), we retained barcodes between 13 and 17 nucleotides in length and excluded all others after extraction. To correct sequencing errors, including insertions and deletions, we applied the Deletion-Correct method^70^. Since samples from each mouse were collected across multiple time points, we pooled all barcode sequences from the same mouse into a single dataset before applying error correction, thereby maintaining consistent barcode identities across time points. This created a master list of consensus barcodes for each mouse, which was then used to track barcode occurrences in individual samples. The final dataset contained read counts for each barcode at each time point, which were used to quantify barcoded lineage dynamics over time.

### Barcode dynamics visualization, diversity and clonal clustering

We used *Doblin*^58^, an R package for barcode lineage analysis, to visualize barcode trajectories, quantify diversity, and cluster dominant lineages based on their temporal dynamics. First, we applied *plot_dynamics()* to visualize barcode frequencies on both logarithmic and linear scales and examine the relative abundance of different lineages over time. To facilitate comparisons of barcode dynamics within and between facilities, we implemented a consistent color-coding scheme, assigning identical colors to identical barcodes detected in multiple mice. To quantify lineage diversity, we used Doblin’s *plot_diversity()* function to calculate the Hill diversity of order 1 (^1^*D*), equivalent to the exponential of Shannon entropy and expressed as the effective number of equally abundant lineages (**Fig. 2a**).

To identify dominant lineage clusters with similar temporal dynamics, we first calculated pairwise similarities between barcode-frequency trajectories using Pearson correlation *r* and used these values to construct a distance (1 - *r*) matrix for clustering. We filtered out low-frequency barcodes, retaining only those with a mean frequency of at least 10^-5^ and detected at a minimum of 11 time points. This filtering reduced the contribution of rare and sparsely observed lineages to the clustering analysis. We then performed hierarchical clustering using the Unweighted Pair Group Method with Arithmetic Mean (UPGMA), implemented in Doblin’s *perform_hierarchical_clustering()* function, to group barcodes based on the similarity of their temporal trajectories. After clustering, we used local regression (LOESS) to generate smoothed representative trajectory for each cluster using *plot_clusters_and_loess.* These smoothed trajectories summarized the temporal behavior of the lineages assigned to each cluster. To determine the selected number of clusters, we used Doblin’s *plot_hc_quantification()* function, which identifies the clustering cutoff to balance cluster separation and frequency ranking in the package (**Extended Data Fig. 2a-b**). This procedure was used to avoid excessive partitioning of lineages with similar temporal trajectories.

### 16S rRNA Profiling

To characterize the bacterial community composition, we amplified the 16S rRNA V4 region from extracted genomic DNA using the 515F/806R-B primer set ^71^. Illumina sequencing adapters were added in a second PCR using the conditions described above for library indexing. The resulting amplicons were pooled, purified, and sequenced on an Illumina MiSeq v2 500-cycle. We used the DADA2 pipeline^72^ to process paired-end Illumina MiSeq reads. After trimming primers and filtering out low-quality reads (Phred score≥ 25), we merged forward and reverse reads and corrected sequencing errors. Chimeras were removed to avoid artifacts. We assigned taxonomy using the SILVA v138 database ^73^ and filtered out any non-bacterial reads like mitochondria and chloroplasts. We used the vegan package ^74^ to rarefy sequencing libraries to a common sampling depth and the Doblin package ^57^ to visualize temporal changes in microbial community composition. Relative abundances were calculated, and family-level diversity (^1^D Hill diversity of order 1) was quantified and visualized with *plot_diversity()*.

### Clustering *E. coli* clonal clusters with community composition

To understand the temporal association between bacterial community and *E. coli* clones, we perform co-clustering as described previously^46^, since these interactions may involve local shifts or temporal delays, For bacterial community dynamics, we analyzed log10-transformed relative abundances of taxa at the family level, including only those present in at least eight time points. For *E. coli* clonal dynamics, we used LOESS-smoothed trajectories derived from barcode clustering. To eliminate amplitude-related effects, all time-series vectors were z-normalized before calculating pairwise shape-based distances. Finally, hierarchical clustering was performed using the average linkage method, generating dendrograms to visualize relationships between lineages and microbial taxa (**Extended Data Fig. 3**).

To determine whether similar temporal patterns occurred across mice, we subsequently extended the analysis to all eight animals (**Extended Data Fig. 6**). *E. coli* lineage-cluster trajectories and family-level bacterial community trajectories from individual mice were combined into a single dataset, with each trajectory retaining its mouse identity. Each trajectory was z-normalized, pairwise SBD values were calculated, and average-linkage hierarchical clustering was performed as described above. This analysis tested whether trajectories from different mice clustered according to temporal similarity irrespective of mouse identity.

### Barcode identity similarity across mice

To determine whether similar cross-mouse lineage dynamics involved the same barcode identities, we calculated the Szymkiewicz-Simpson overlap coefficient^75^, which quantifies the overlap between the barcode sets assigned to two lineage clusters relative to the size of the smaller set. The coefficient ranges from 0 (no overlap) to 1 (complete containment of the smaller set within the larger, or identical when both are of equal size) and was used to quantify shared barcode identity between clusters from different mice. To determine whether the observed overlap exceeded that expected by chance, we generated null barcode sets by randomly sampling from the full barcode pool while preserving the observed sizes of the two clusters, and recalculated the overlap coefficient for each simulated pair. We repeated this procedure 1,000 times to generate a null distribution of expected overlap coefficients. The significance of each observed barcode overlap was then assessed using a z-score calculated relative to the mean and standard deviation of the corresponding null distribution. Clusters with z > 1.96 were considered to show greater-than-expected barcode overlap at the nominal 5% significance level.

### Dynamic Covariance Mapping of community interactions

We used Dynamic Covariance Mapping (DCM)^46^ to infer time-progressive community interaction matrices for each mouse from the combined *E. coli* lineage-cluster and bacterial family-level time series. In DCM, the community state at time *t* is represented by the vector *x*(*t*) = (*x*_1_(*t*), …, *xₙ*(*t*)*^T^, where each element denotes the processed trajectory of an *E. coli* lineage cluster or bacterial family. Following DCM-specific preprocessing of the individual trajectories, community dynamics were represented as *x^•^* = *f*(*x*), where *f*(*x*) defines the temporal rates of change of the community state variables. Although the full nonlinear dynamics described by *f*(*x*) are unknown ^8,10,76,77^, local system behavior around a reference state *x*^∗^ = *x*(*t*^∗^) can be approximated to first order by a Taylor expansion: *x^•^* ≈ *f*(*x*^∗^) + *J*^∗^(*x* − *x*^∗^), where *J*^∗^ is the local community Jacobian, with elements *J_i_*_j_ = *ϱf_i_*/*ϱx*_j_|*_x_*_=*x*∗_. Under this local linear approximation, the cross-covariance between the rates of change and community states is related to the Jacobian by, Cov(*x^•^*, *x*) ≈ *J*^∗^Σ*_x_*, where Σ*_x_* = Cov(*x*, *x*) is the covariance matrix of the community state variables within the estimation window. Following the DCM framework^46^, the entries of Cov(*x^•^*, *x*) were used directly as covariance-based estimates of community interaction structure. For the primary analysis, within each 5-step sliding time window *τ_k_*, the DCM interaction estimate was calculated as Cov*_τk_*(*x^•^_i_*, *x*_j_). The ordering of the indices specifies the direction represented by the DCM estimate *j* → *i*: its sign distinguishes positive from negative inferred interactions, whereas its magnitude quantifies the strength of the corresponding dynamic covariance over interval *τ_k_*. Sensitivity to temporal-window choice was assessed by repeating the analysis using sliding windows spanning 1 to 10 sampling steps.

We generated a sequence of DCM interaction matrices over successive sliding time windows and calculated the eigenvalues of each matrix. The full eigenvalue spectrum was examined to characterize temporal changes in community dynamics. For each matrix, the spectral abscissa, *max_i_ Re(λ_l,τk_*, denoted as Re(*λ*_max_) and defined as the largest real part among the eigenvalues, was used as a DCM-based summary of local dynamical convergence or divergence over time. For a system linearized around an equilibrium, negative values indicate local asymptotic stability, positive values indicate at least one locally divergent direction, and zero defines the marginal-stability boundary. Because the sampled communities were not assumed to be at equilibrium, we interpreted the spectral abscissa more generally as an indicator of local dynamical convergence or divergence rather than as a formal test of equilibrium stability. To quantify temporal changes in interaction estimates, we calculated the difference in each DCM interaction estimate between successive sliding windows, *Δĵ_i,j,k_ = ĵ^DCM^_ij,τk_ − ĵ^DCM^_ij,τk−1_*, and quantified the magnitude of change as |Δĵ_i,j,k_|. Interaction elements showing a concordant direction of change in at least five of the eight mice were retained for further analysis. For these recurrent interactions, we tracked their values across successive windows and quantified similarities among their temporal trajectories using dynamic time warping (DTW). The resulting DTW distance matrix was subsequently used to cluster interactions according to their temporal behavior.

### Isolation of Paenibacillaceae from fecal samples

To isolate Paenibacillaceae from fecal samples, we thawed samples at 3, 6, and 12 h post-gavage and on day 12 from mice m5, m6, m7, and m8 at room temperature for 10 minutes, followed by brief centrifugation for 30 s. Because members of Paenibacillaceae are capable of forming heat-resistant endospores, we used heat treatment to selectively eliminate non-spore-forming microbes. A 10 µL aliquot of fecal suspension was added to 500 µL of sterile water and incubated at 80°C for 10 min. This step ensured that only spore forming Paenibacillaceae survived, preventing them from being outcompeted by barcoded *E. coli* if plated immediately. After heat treatment, 100 µL of each day-12 sample was plated onto LB agar containing spectinomycin (100ug/ml), whereas samples from earlier time points (3-12h) were plated onto LB agar without antibiotic. The plates were placed in a box with AnaeroPack at 30°.After 48 hours, we observed visible colonies on plates for day 12 while no colonies were detected from the earlier time points. To confirm that the isolates were indeed Paenibacillaceae, we extracted genomic DNA from selected colonies and amplified the 16S rRNA region using Sanger sequencing. The sequencing results confirmed that the isolated colonies belonged to the Paenibacillaceae family.

### Whole-genome sequencing, assembly, and resistance-mechanism analysis of Paenibacillaceae isolates

Glycerol stocks of the *Paenibacillus* isolates recovered from day-12 of mice m5–m8 were streaked onto LB agar with spectinomycin (100 µg/ml) and grown anaerobically (AnaeroPack) at 30 °C. A single colony per isolate was expanded in LB–spectinomycin broth, and the cell pellet was submitted for Oxford Nanopore long-read whole-genome sequencing (R10.4.1 flow cells; super-accurate basecalling, Dorado v4.3.0). Genomes assembled by PlasmiSaurus long-read pipeline: the lowest-quality 5% of reads were removed with Filtlong v0.2.1, downsampled to 250 Mb, and a draft sketch generated with Miniasm v0.3 to estimate genome size and coverage; reads were then re-downsampled to ∼100× with strong quality weighting, assembled de novo with Flye v2.9.1, and polished with Medaka v1.8.0. Completeness and contamination were assessed with CheckM2, and genomes were annotated with Bakta v1.12.0 (database v6.0). Three isolates (m5, m6, m8) assembled into single circular chromosomes (∼7.47 Mb; 84–103× coverage; 99.85% complete, 0.06–0.12% contamination); the m7 isolate (9×, seven contigs) was excluded from downstream analyses. Species identity was confirmed by FastANI v1.34 against the complete *Paenibacillus macerans* I6 reference genome (ATCC 7068; GCF_022494515.1), giving 99.26% average nucleotide identity; pairwise comparisons showed the isolates to be effectively clonal.

To test whether resistance was acquired horizontally from the colonising E. coli, we searched each assembly for the *E. coli* spectinomycin-resistance cassette (*aadA*) by BLASTn and screened all genomes for any annotated antimicrobial-resistance determinant including the Gram-positive spectinomycin adenylyltransferases *aad9* using AMRFinderPlus, the Resistance Gene Identifier (RGI; CARD), and ResFinder. Whole-genome structure was compared with nucmer (MUMmer v4.0.1) and a pan-genome was built with Panaroo v1.5.2; all isolate-specific regions and accessory gene families were annotated with Bakta and screened for resistance cargo. To identify candidate resistance mutations, we extracted the two components of the spectinomycin binding site, 16S rRNA helix 34 and ribosomal protein S5 (*rpsE*) from each isolate and aligned them to I6 and to spectinomycin-sensitive reference taxa (*E. coli* K-12, B. subtilis 168) with MAFFT v7.526. The *rpsE* variant was validated at the read level by mapping isolate reads to the I6 rpsE locus with minimap2 v2.30 (map-ont) and samtools v1.23.1.

## Data availability

Raw barcode sequencing data from this study have been deposited in the National Center for Biotechnology Information Sequence Read Archive. This includes BioProject accession number <u>PRJNA1513701</u> for all 16S data, <u>PRJNA1513734</u> for all high-resolution barcode data from, and <u>PRJNA1514206</u> for whole genome sequencing results of *Paenibacillus macerans* isolates.

## Code availability

Code is available via GitHub at https://github.com/melisgncl/high-resolution-gut-microbiome-dynamics-during-antibiotic-perturbation/tree/master.

## Acknowledgments

We thank the mice facilities at Université Sherbrooke. This work was supported by the following grants: Canadian Institute for Health Research PG-408523 (AS), Natural Sciences and Engineering Research Council of Canada RGPIN-2016-06566 (AS), Canada Research Chairs (AS), National Research Foundation of South Africa 89967 (CH), and Human Frontier Science Program RGP027/2026 (CH).

## Author contributions

AS and MG conceptualized this study. GMC, AF, and SR performed the *E. coli* colonization experiments. MG performed the genomic extraction, barcode amplification, 16S rRNA profiling with the help of CM. MG and AS implemented all the bioinformatic analysis. AS, MG, and CH developed the dynamic covariance mapping approach and ecological analyses. AS and MG wrote the manuscript with feedback and editorial support from all authors.

## Extended Data Figures

**Extended Data Figure 1:**
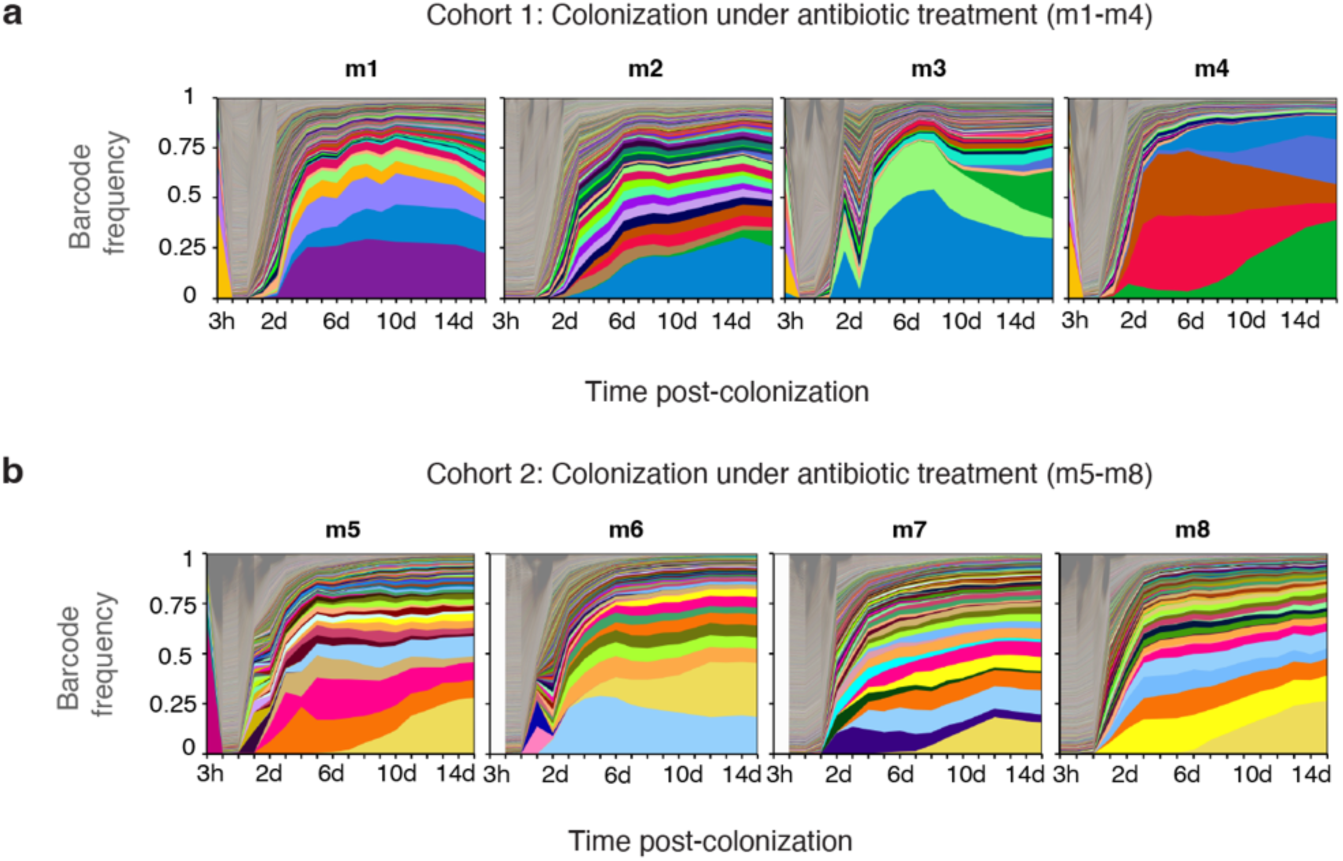
Barcode dynamics and diversity in colonized mice (m1–m8), related to. Fig. 1 **a-b,** Barcode frequency dynamics for cohort 1 (m1–m4) and cohort 2 (m5–m8), shown on a linear scale. Each panel represents a single mouse (m1 through m8). Same colors across different mice correspond to identical chromosomal barcodes.

**Extended Data Figure 2:**
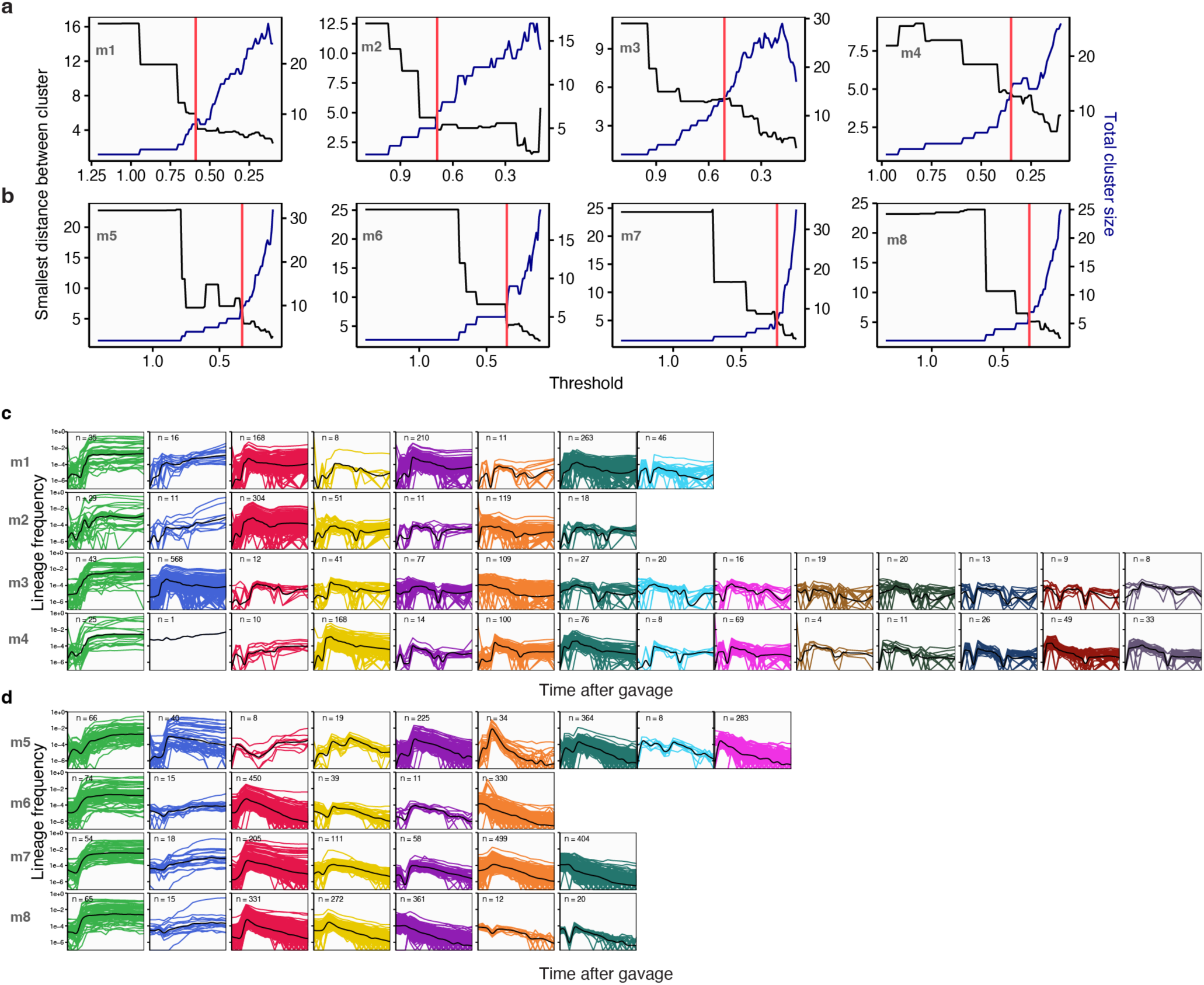
Identification of dominant *E. coli* clonal clusters, related to. Fig. 1 **(a-b)** Dominant *E. coli* clonal clusters were identified using hierarchical clustering implemented in the Doblin package. To determine the optimal number of clusters, we used Doblin heuristic method, which compares the Euclidean distance between consensus trajectories (black curve) and the number of resulting clusters (blue curve). The intersection point (red line) indicates the optimal clustering cutoff. This threshold was used to define the final dominant clonal clusters used in downstream analyses. Panel a shows results for cohort 1 (m1–m4), and b for cohort 2 (m5– m8), with each panel representing one mouse. **(c)** Dominant *E. coli* clonal clusters in cohort 1. Each colored line represents the time-series trajectory of a unique barcode within a cluster; color assignments match those in Figure 2a. Black lines show the LOESS-smoothed consensus trajectory for each cluster. The number of raw barcodes per cluster is indicated above each panel. Colors correspond to those used in Figure 1e-f. **(d)** Same as (c), but for cohort 2 (m5–m8).

**Extended Data Figure 3:**
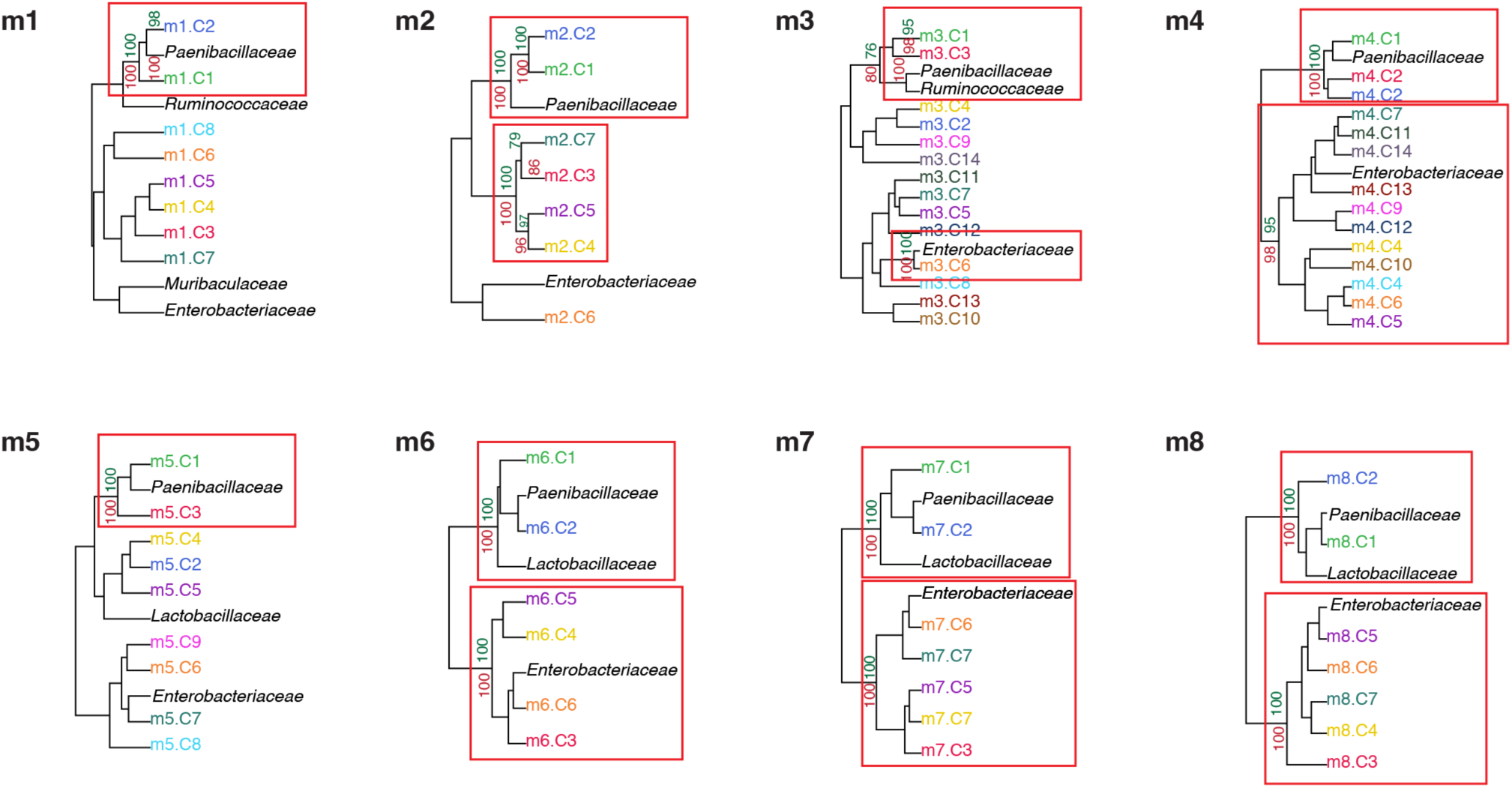
Statistical support for clustering of *E. coli* clonal clusters with bacterial family composition, related to. Fig. 1 *pvclust* was used to assess the robustness of hierarchical clustering in cohort 1 (m1–m4, top row) and cohort 2 (m5–m8, bottom row). These co-clustering results correspond to those shown in Figure 1e-f. We performed 1,000 bootstrap resampling runs to evaluate cluster stability. In each dendrogram, approximately unbiased (au) scores are shown in red, and bootstrap probabilities are shown in green on the left. Clusters with au≥ 95 are highlighted with rectangles, indicating strong statistical support. Across all mice, clusters linking Paenibacillaceae with either clonal cluster C1 or C2 consistently exhibited high au scores and bootstrap probabilities.

**Extended Data Figure 4.**
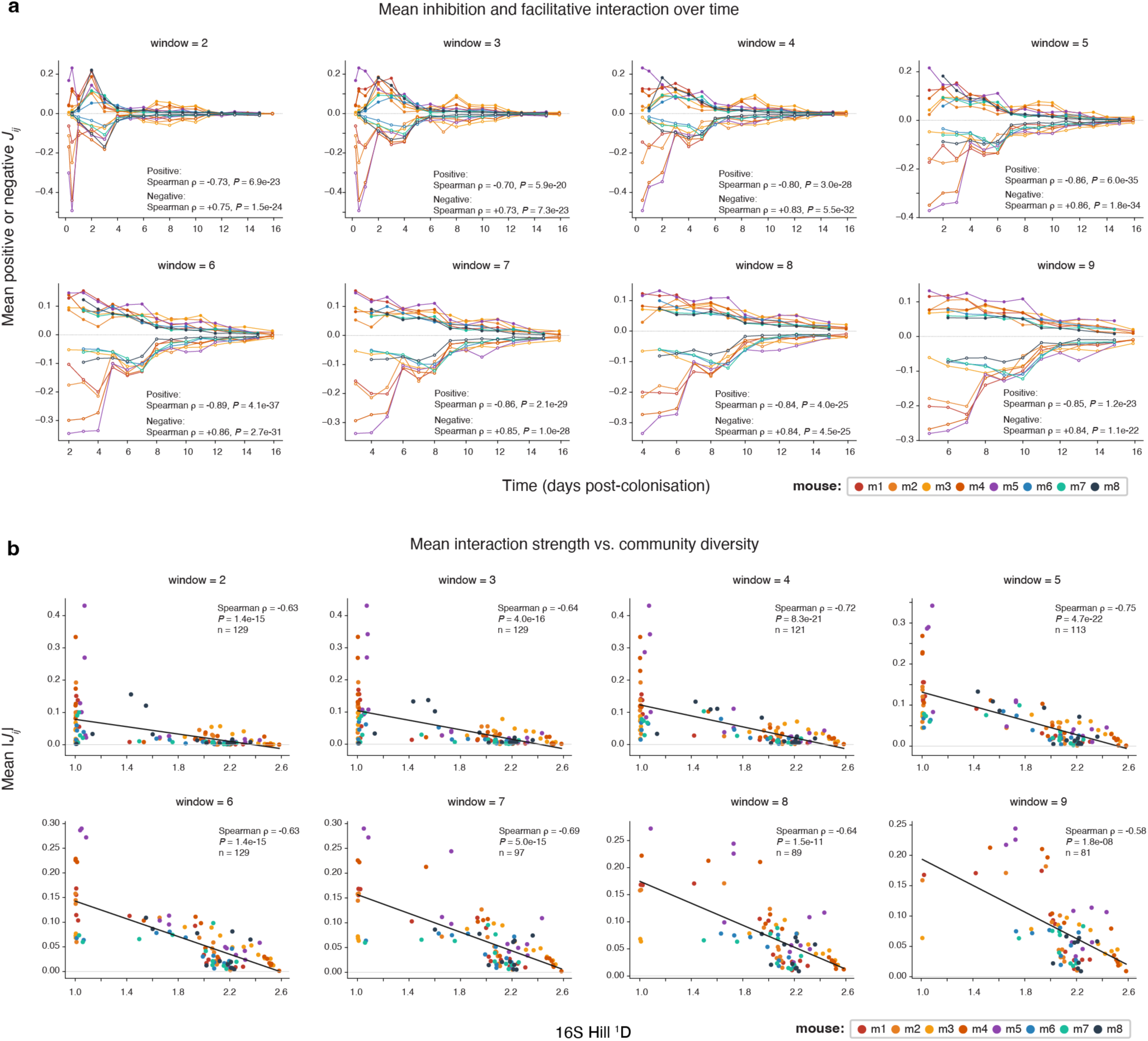
Window-choice robustness of the community interaction strength and diversity trends. **(a)** Mean positive (filled markers) and mean negative (open markers, plotted as magnitude) *J*ᵢⱼ versus time post-colonisation, one panel per sliding-window size (window = 2–9), pooled across all eight colonised mice (colours as in Fig. 2). Both branches decline toward zero with time at every window tested (Spearman ρ range: positive, −0.70 to −0.89; negative magnitude, −0.73 to −0.86; all *P* < 10⁻¹⁹). The primary window (window = 5, used throughout the main text) gives ρ = −0.86 for both branches (*P* = 6.0×10⁻³⁵ and 1.8×10⁻³⁴, n = 113), reproducing Fig. 2c exactly. **(b)** Mean absolute interaction strength |*J*ᵢⱼ| versus community diversity (16S Hill ¹D), one panel per sliding-window size (window = 2–9), pooled across mice and sampled days. The negative relationship between interaction strength and diversity holds at every window (Spearman ρ range: −0.58 to −0.75; Pearson r shown for reference), strongest around window = 5–6 and weakening somewhat by window = 9 as sample size drops (n = 129 at window 2 to n = 81 at window 9, reflecting fewer valid sliding windows as window size grows). Window = 5 gives ρ = −0.75 (*P* = 4.7×10⁻²², n = 113), reproducing Fig. 2d exactly. Together, both panels show that neither trend reported in the main text is an artefact of the window = 5 choice, both hold across the full range of windows tested (2– 9), with window = 5 sitting near the point of strongest signal in each case.

**Extended Data Figure 5.**
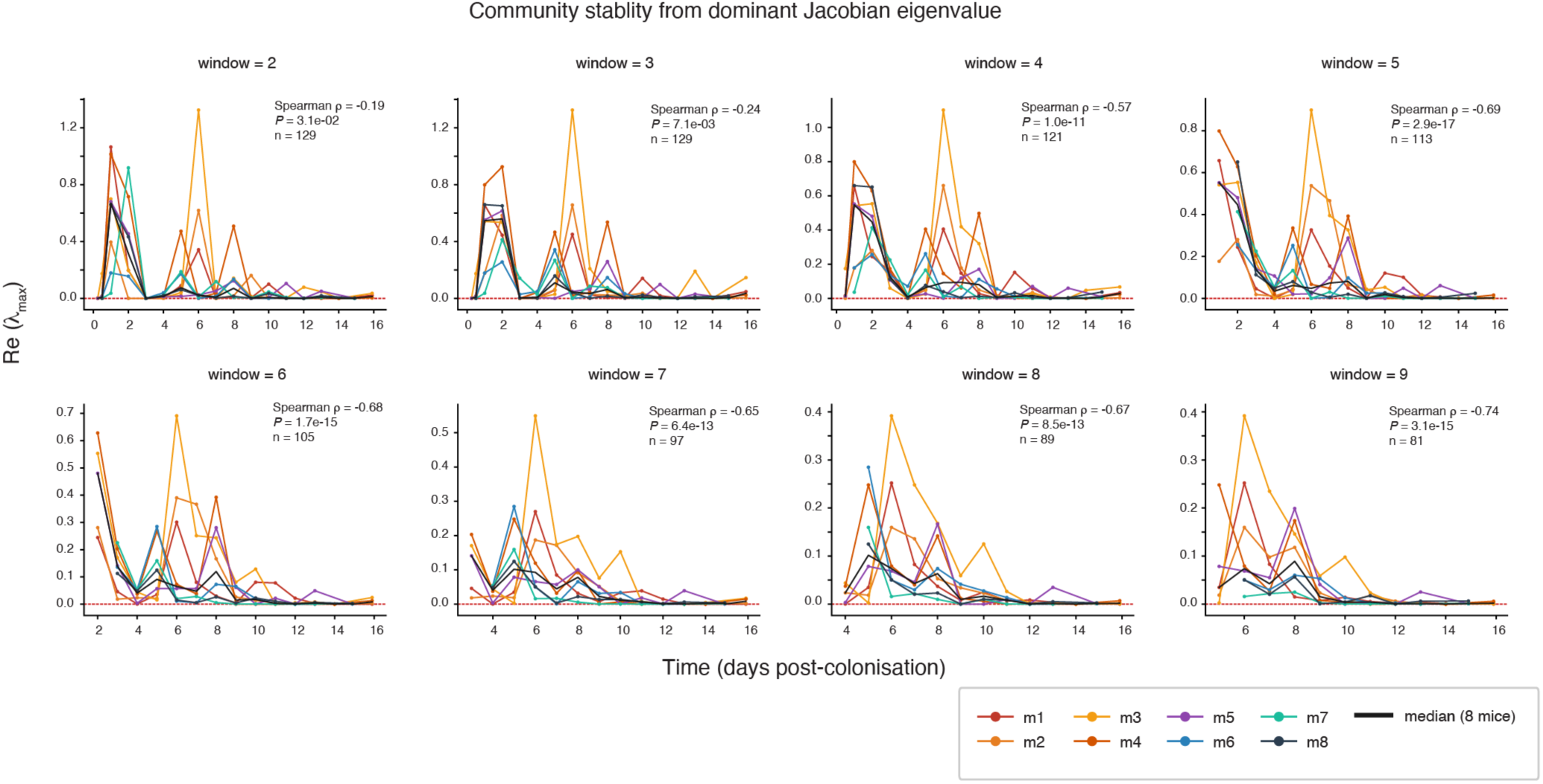
Community stability as a function of window sizes. Dominant Jacobian eigenvalue Re(*λ_max_*) versus time post-colonisation, one panel per sliding-window size (window = 2–9), pooled across all eight colonised mice (colours as in Fig. 3; black line, cross-mouse median). Unlike the interaction-strength trends in Fig. 2c-d, this relationship is window-dependent: at window = 2–3 the decline is only weakly monotonic (Spearman *ρ* = −0.19, *P* = 0.031, and ρ = −0.24, *P* = 0.007) because a secondary bump near day 5–6, coincident with Paenibacillaceae onset and driven largely by m3, interrupts the early decline. From window = 4 onward this bump is absorbed into a smoother decline and ρ stabilises between −0.57 and −0.74 (all *P* < 10⁻¹¹). The primary window (window = 5, used throughout the main text) gives ρ = −0.69 (*P* = 2.89×10⁻¹⁷, n = 113), reproducing Fig. 3a exactly, at a point where the trend has already stabilised rather than at its strongest.

**Extended Data Figure 6:**
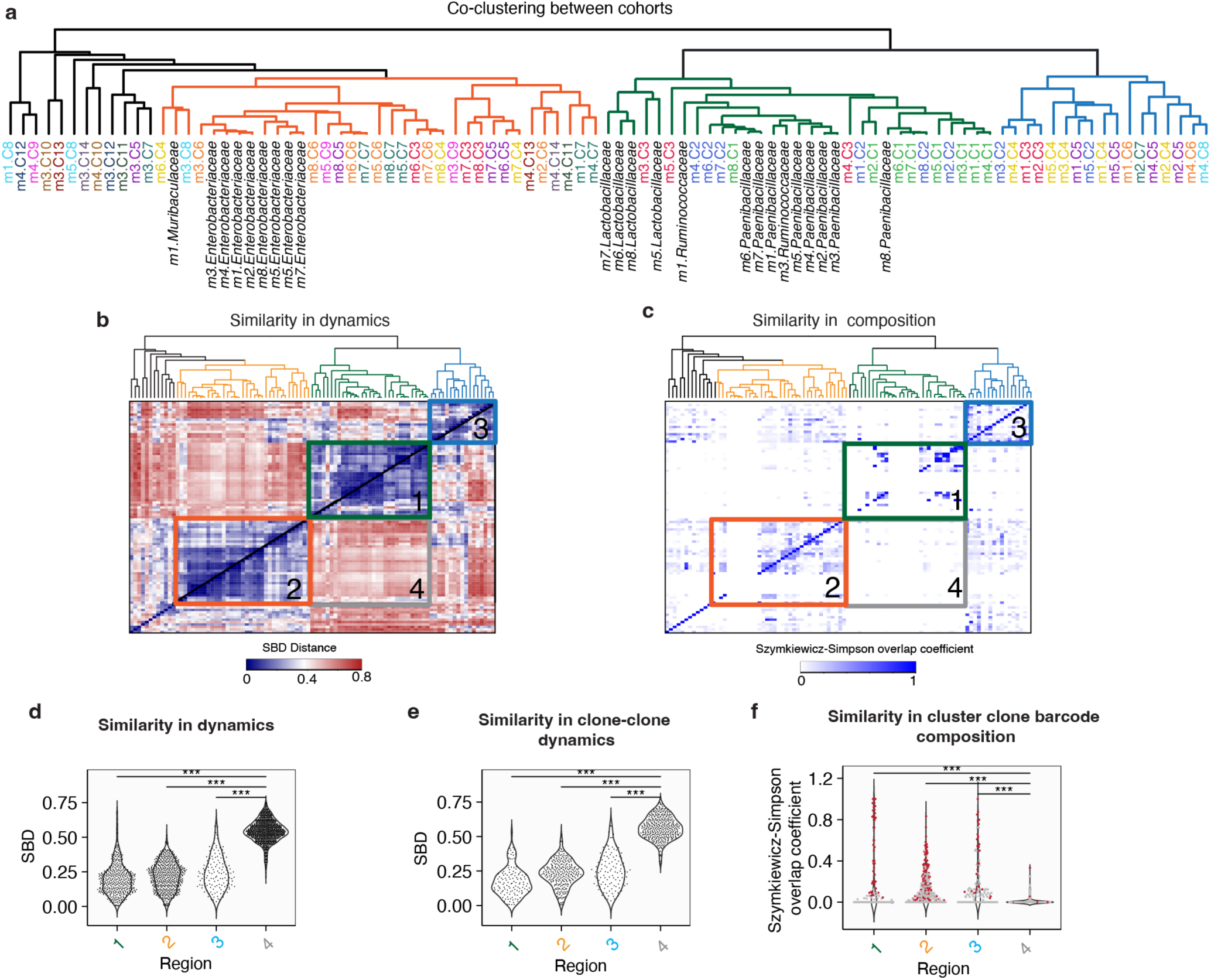
Temporal dynamics and barcode composition comparisons within cohorts 1 and 2, related to Fig. 5. **(a)** Dendrogram generated from co-clustering analysis of barcode and bacterial family abundance trajectories in cohorts 1 and 2, using shape-based distance (SBD) metrics. **(b)** Heatmap of the pairwise SBD matrix for all clone–family pairs. Rows and columns are ordered according to the dendrogram in panel **a**. Based on dynamic similarity, three distinct regions were identified: Region 1 (green), comprising Paenibacillaceae families and high-frequency clonal clusters (C1 and C2); Region 2 (orange), consisting of Enterobacteriaceae-associated families and low-frequency clones; and Region 3 (blue), containing transient clusters that exhibited early abundance increases followed by plateauing. (Lower SBD values indicate greater similarity in temporal dynamics.) **(c)** Matrix of Szymkiewicz–Simpson overlap coefficients showing barcode identity overlap between clonal clusters. Matrix layout and clustering match those in panel **b**. **(d)** Violin plots displaying the distribution of pairwise SBD values within and between regions. Region 4 serves as a cross-region control. Intra-region clone or family pairs exhibited significantly more similar dynamics than inter-region pairs (\*\*\**P* < 0.001, Wilcoxon test). **(e)** Same analysis as in panel **d** but restricted to clone–clone SBD comparisons. **(f)** Szymkiewicz–Simpson overlap coefficients for barcode identity in clone–clone pairs within Regions 1, 2, and 3, compared to cross-region pairs (Region 4). Statistically significant overlaps, determined via bootstrapping (see Methods), are highlighted in red.

